# Loss of MAGEL2 Disrupts Pituitary Translation in a Mouse Model of PWS and Schaaf–Yang Syndrome

**DOI:** 10.64898/2026.05.12.724462

**Authors:** Tara Bayat, Maria Camila Hoyos Sanchez, Cristian Camilo Rodríguez-Almonacid, Denis Štepihar, Elena B Tikhonova, Farzana Yeasmin Popy, Juan Sebastian Solano Gutierrez, Stephanie Myers, Miloš Vittori, Andrey L. Karamyshev, Zemfira N. Karamysheva, Klementina Fon Tacer

**Affiliations:** School of Veterinary Medicine, Texas Tech University, Amarillo, TX 79106, USA; Texas Center for Comparative Cancer Research (TC3R), Amarillo, TX 79106, USA; Department of Cell Biology and Biochemistry, Texas Tech University Health Sciences Center, Lubbock, TX 79430, USA; Center for Biotechnology and Genomic Medicine, Medical College of Georgia, Augusta University, Augusta, GA 30912, USA; University of Ljubljana, Biotechnical Faculty, Department of Biology, Ljubljana, Slovenia

**Keywords:** MAGEL2, Prader-Willi Syndrome, Schaaf-Yang Syndrome, polysome profiling, pituitary, translation, TEM, secretory granules

## Abstract

Prader-Willi syndrome (PWS) and Schaaf-Yang syndrome (SYS) are neurodevelopmental disorders associated with hypothalamic-pituitary dysregulation. In the pituitary gland, translational control enables rapid peptide hormone production and secretion in response to hypothalamic signals without requiring new mRNA synthesis, yet the mechanisms regulating pituitary translation remain poorly understood. Furthermore, although the PWS-associated gene *MAGEL2* has been implicated in neuroendocrine regulation and vesicular trafficking in the hypothalamus, its role in the pituitary gland remains unknown. Initial analysis of previously published pituitary proteomic data revealed enrichment of translation-associated pathways among downregulated proteins in *Magel2* KO mice, suggesting translational impairment. Here, we investigated the impact of *Magel2* loss on pituitary translatome using polysome profiling and RNA sequencing. We first optimized a polysome profiling workflow for mouse pituitary tissue and established that pooling two to three pituitaries yielded sufficient RNA quality and quantity for downstream analyses. Polysome profiling of WT and *Magel2* KO pituitaries revealed no major alterations in global translational activity, as translated and nontranslated fractions were largely unchanged between genotypes. However, transmission electron microscopy revealed a shift toward smaller secretory granule size, indicating altered granule maturation dynamics. To further characterize the pituitary translatome, RNA sequencing was performed on input, monosome, light polysome, and heavy polysome fractions. Clustering analyses identified six distinct translational trajectories across fractions, revealing fraction-specific enrichment of biological pathways. RNAs enriched in heavy polysomes were associated with metabolic and oxidative phosphorylation pathways, whereas monosome-enriched clusters were linked to RNA processing and translation-related functions, suggesting specialized translational regulation within the pituitary. Differential expression analysis demonstrated that translatomic alterations were more pronounced than transcriptomic changes in *Magel2* KO pituitaries, with the strongest enrichment observed in heavy polysome fractions. Functional enrichment analyses identified pathways associated with endocrine and metabolic regulation, circadian rhythm, cytoskeleton organization, vesicular trafficking, and RNA regulation, suggesting that translation contributes to pituitary physiological function and patient symptoms. For example, prolactin displayed altered polysome association without changes at the transcript level, consistent with the increased serum prolactin levels observed in *Magel2* KO mice and in patients with PWS. Interestingly, the PWS-associated gene *Necdin* (*Ndn*) was consistently downregulated across all fractions, which contrasts with previously described compensatory upregulation in the hypothalamus. Together, our findings suggest the involvement of MAGEL2 in pituitary in transcriptional and translational processes and the organization of the secretory pathway and provide the first comprehensive characterization of the mouse pituitary translatome. This work provides new insights into the mechanisms underlying neuroendocrine dysfunction in PWS and SYS and establishes a resource for future studies of translational regulation in neuroendocrine disease.

## INTRODUCTION

Gene expression is regulated at multiple levels, with translational control serving as a central mechanism that enables rapid and precise adjustment of protein synthesis in response to physiological and pathological conditions. This layer of regulation accounts for a substantial portion of protein variability and plays critical roles in processes such as neural plasticity, endocrine function, and reproductive signaling. Unlike transcriptional regulation, translational control allows cells to rapidly adjust protein levels without altering mRNA abundance, enabling precise and timely responses to environmental changes, stress, nutrient availability, and disease states. As such, translational control is a key mechanism for maintaining cellular homeostasis and adapting to dynamic biological demands. Notably, nearly half of the variation in protein concentration has been attributed to translational regulation, underscoring its central role in fine-tuning gene expression and cellular function [1–5].

In highly active endocrine cells, such as those in the hypothalamus and pituitary, robust protein translation is essential for hormone production and efficient regulation of physiological functions, including reproduction. This is supported by the discordance between the pituitary transcriptome and proteome and the identification of ribosomal proteins as markers of pubertal development [6–8]. Furthermore, pathological mutations affecting ribosome dynamics and translation efficiency have been shown to contribute to disorders such as neurodegeneration, cystic fibrosis, and autism, as well as genetic diseases including Diamond–Blackfan anemia and Treacher Collins syndrome [9–20]. However, very little is known about how translation is regulated in these tissues, and its role in Prader-Willi syndrome (PWS), Schaaf-Yang syndrome (SYS), and related neurodevelopmental disorders remains largely unexplored.

PWS and SYS are complex imprinted neurodevelopmental disorders, caused by loss of one or more paternally inherited genes located on the human 15q11-q13 locus [21, 22]. One of the protein-coding genes affected by paternal deletion in PWS is melanoma antigen L2 (*MAGEL2*), which also harbors point mutations in SYS [21, 23, 24]. Individuals with PWS and SYS share several clinical features, including hypotonia, hyperphagia, autism spectrum disorder, intellectual disability, neurodevelopmental delay, sleep disturbances, and neuroendocrine dysfunction, suggesting an important contribution of MAGEL2 to these phenotypes [25]. Furthermore, many features of these symptoms are recapitulated in mouse models with depleted *Magel2* gene [25] and linked to hypothalamic abnormalities and impaired pituitary endocrine function [13, 14, 21, 26–38]. Recent proteomic analysis of wild-type and *Magel2* knockout (KO) pituitaries identified differential expression of ribosomal and signal recognition particle (SRP) proteins, suggesting a role for MAGEL2 in mRNA translation in addition to well established protein ubiquitination and recycling [39–43]. Given that pituitary hormones such as prolactin are secreted peptide hormones, efficient signal recognition particle (SRP)-mediated cotranslational targeting of nascent proteins to the endoplasmic reticulum is essential for hormone biosynthesis and secretion in highly specialized pituitary endocrine cells. SRPs recognize N-terminal signal sequences of secretory proteins and direct ribosome–nascent chain complexes to the ER membrane for protein translocation. Importantly, inefficient SRP interaction with nascent chains can trigger an mRNA quality control pathway, and SRP depletion has been shown to reduce levels of secretory proteins [44]. Therefore, MAGEL2 may impact multiple aspects of the translational and secretory pathways, in addition to its previously described role in endosomal protein recycling.

MAGEL2 was shown to function at endosomes as part of the MUST complex (MAGEL2-USP7-TRIM27), comprising the E3 ubiquitin ligase TRIM27 and the deubiquitinase USP7. This complex regulates ubiquitination of the WASH complex to control F-actin nucleation, thereby enabling retromer-dependent protein recycling [25, 40, 45, 46]. Preliminary proteomic data in *Magel2* KO mice, along with recent findings of upregulated ribosomal protein genes in post-mortem brain and iPSC-derived neural progenitor cells from individuals with autism, suggest that MAGEL2 may impact translation [39, 47]. Given that MAGEL2 is highly expressed in the hypothalamus in both humans and mice, as well as in the human pituitary [21], we hypothesized that it plays an important role in multiple aspects of hormone production and secretion, including translational regulation. In this study we aimed to investigate, for the first time, the role of MAGEL2 in pituitary mRNA translation using polysome profiling.

Polysome profiling is a widely used approach to study mRNA translation by assessing the association of transcripts with ribosomes, thereby defining the translatome, the subset of mRNAs actively translated within a cell. This method relies on sucrose gradient fractionation to separate untranslated mRNAs from those associated with monosomes or polysomes, reflecting increasing levels of translational engagement. Compared with other techniques, polysome profiling provides a robust measure of translational activity by quantifying ribosome occupancy on individual transcripts. It is suitable for both targeted analyses and global assessments of translational efficiency [1, 2, 48–50]. However, it has not been widely applied to highly active and secretory endocrine tissues such as the pituitary gland due to technical challenges, including the limited tissue availability. This often necessitates pooling of samples. In this study, we addressed these limitations by optimizing and successfully applying polysome profiling to murine pituitary tissues, enabling the first in-depth analysis of translation in pituitary, particularly in the context of neurodevelopmental syndromes involving endocrine dysfunction, such as PWS and SYS. Our results provide a resource for the pituitary translatome and establish a foundation for investigating translational regulation in the pituitary, including potential novel roles of MAGEL2 in gene expression and protein synthesis and the contribution of translational dysregulation to PWS and SYS.

## MATERIALS AND METHODS

### Mouse colony housing, breeding strategy, and genotyping

All animal procedures were performed in accordance with protocols approved by the Institutional Animal Care and Use Committee (IACUC) at Texas Tech University. C57BL/6-*Magel2*^tm1Stw/^J mice (stock no. 009062) and C57BL/6 wild-type (WT; stock no. 000664, B6) controls were originally obtained from The Jackson Laboratory (Bar Harbor, ME, USA). The *Magel2*^tm1Stw^ mouse carries a maternally inherited imprinted WT allele and a paternally inherited *Magel2*-lacZ knock-in/knockout allele, resulting in loss of endogenous *Magel2* expression (*Magel2*^pΔ/m+^), hereafter referred to as *Magel2* knockout (KO). On a C57BL/6J background, these mice exhibit several features of Prader–Willi syndrome (PWS) and Schaaf-Yang syndrome (SYS) and are widely used as a PWS/SYS model [21, 51–54]. Mice were housed under standard laboratory conditions with corncob bedding on a 12-h light/dark cycle (lights on at 06:00 and off at 18:00), at an ambient temperature of 23–24°C and relative humidity of 30–35%. Animals had *ad libitum* access to irradiated, extruded, soy-free rodent chow (2916, Inotiv) and filtered water.

For colony maintenance and generation of experimental cohorts (Figure S1A), heterozygous females (*Magel2*^p+/Δm^) were mated with C57BL/6 WT males. These females carry a deletion on the maternally inherited allele, which is imprinted and transcriptionally silenced. Consequently, offspring inheriting the maternal *Magel2* KO allele (*Magel2*^p+/Δm^) are phenotypically indistinguishable from WT littermates. Both male and female heterozygous mice generated using this breeding scheme carry the mutation on the maternal allele only and were therefore used as phenotypically WT controls. To generate phenotypically *Magel2* KO mice for experiments, heterozygous males (*Magel2*^pΔ/m+^) were bred with C57BL/6 WT females. This cross yields offspring with a paternally inherited *Magel2* deletion (*Magel2*^pΔ/m+^), resulting in functional KO animals due to silencing of the maternal allele by imprinting.

Pups were weaned at postnatal day 21, and tail biopsies were collected for genotyping. Genomic DNA was extracted using the NaOH method as previously described [55]. Genotyping was performed by PCR using the KAPA2G DNA polymerase kit (Sigma-Aldrich), followed by electrophoresis on a 2% agarose gel (Figure S1B). The following primers were used (Jackson Laboratory protocol): forward primer 1, GAT GGA AAG ACC CTT GAG GT; forward primer 2, ATG GCT CCA TCA GGA GAA C; and reverse primer, GGG ATA GGT CAC GTT GGT GT. Expected PCR product sizes were 233 bp for the WT allele and 366 bp for the *Magel2* KO allele.

### Tissue collection for polysome profiling optimization and experiments

Polysome profiling optimization was performed using tissues from 7-month-old WT male mice. For the final experiment, tissues were collected from eight WT and eight *Magel2* KO male mice aged 8-10 weeks. Mice were euthanized at 9:00 AM by isoflurane inhalation, followed by exsanguination via cardiac puncture. Body and tissue weights were recorded (Figure S1C), and blood was collected for serum preparation. Blood samples were allowed to clot at room temperature for 60 minutes and then centrifuged at 2,000 × g for 15 minutes at 4°C. Serum was collected and stored at −80°C until further analysis. Tissues were rapidly dissected, snap-frozen in liquid nitrogen, and stored at −80°C until use for polysome profiling.

### Mice perfusion and tissue preparation for transmission electron microscopy

Transmission electron microscopy (TEM) was performed using tissues from perfused 6-month-old female WT and *Magel2* KO mice. At 10:00 AM, mice were anesthetized with isoflurane and exsanguinated via cardiac puncture, followed by brief perfusion with PBS containing 10 U/mL heparin and fixation by perfusion with a solution of 2.5% glutaraldehyde and 2% paraformaldehyde in PBS. Whole brains were dissected and stored in the same fixative at 4°C until further processing. For preparation of coronal brain sections, fixative was washed out with PBS, and brains were sectioned using a coronal mouse brain matrix with 0.5-mm spacing (Ted Pella, Inc.) and double-edged blades. Sequential sections were transferred to 24-well plates containing PBS and examined under a Stemi 508 stereomicroscope (Zeiss) to identify sections containing the hypothalamus. Selected sections were stored in 2.5% glutaraldehyde and 2% paraformaldehyde in PBS until embedding.

Samples were then processed in 2.5% glutaraldehyde prepared in 0.05M sodium cacodylate buffer (pH 7.2-7.4). Then, they were rinsed in cacodylate buffer and post fixed in 1% osmium tetroxide. Following additional buffer washes, samples were dehydrated though an ethanol series and transitioned through acetone before infiltration with epoxy resin (Epon/LX-112 equivalent). Samples were embedded and polymerized for 48 hours in 80 °C. Glass knives were created using the LBK Knife Maker 7800 B and mounted onto the RMC PowerTome Ultramicrotome to collect semithin sections stained with methylene blue-azure II and viewed on Olympus BX40 light microscope. Ultrathin sections were obtained with a diamond knife. Sections were collected onto electron microscopy sciences 200 mesh copper grids. Sections were positively contrasted with uranyl acetate and Reynolds lead citrate prior to imaging on the Hitachi H-7650 transmission electron microscope.

### Analysis of TEM images for secretory granule quantification

Contrasted ultrathin sections were collected by Hitachi H-7650 transmission electron microscope (Hitachi Hight - Tech Corp), equipped with AMT 600 camera. Image analysis was performed using ImageJ (version 2.16.0/1.54p). Cells from two biological replicates were analyzed. Secretory granules (SGs) were defined as electron-dense, approximately spherical structures with diameters ranging from 100 to 800 nm and a clearly distinguishable boundary from the surrounding cytoplasm [56]. For each image, the scale was calibrated using the embedded scale bar. All SGs were counted, and their diameters were measured (n = 20 WT images; n = 10 *Magel2* KO images). The total analyzed area per image was determined and used to calculate SG density (number of SGs per μm²). In addition, the mean SG area (μm²) was calculated for each image and used to compare granule size between groups.

### Polysome profiling

Cell lysates preparation and polysome profiling was prepared according to the protocol described previously [2]. Briefly, tissues were lysed in 900 µL of lysis buffer (20 mM HEPES-KOH, pH 7.5, 10 mM MgCl2, 100 mM KCl, 2 mM DTT, 1% NP-40, 1X protease inhibitor cocktail (EDTA-free), 80 units/mL RNasin, and 100 µg/mL of cycloheximide) by mechanical disruption (10 strokes in glass Dounce homogenizer, followed by 10 strokes with a syringe). The lysates were then centrifugated at 12,000 x g for 8 minutes at 4°C, and the clarified supernatant loaded onto a sucrose gradient for polysome separation. 10% of the lysate was used for RNA preparation and analyzed as input material (total transcriptome).

For polysome separation, 500 µL of clarified lysate was loaded on top of 10-50% sucrose gradient prepared in the lysis buffer (20 mM HEPES-KOH (pH 7.4), 10 mM MgCl2, 100 mM KCl, 1 mM DTT and 1X protease inhibitor cocktail). The gradient tubes were ultracentrifuged at 260,000 x g for 2 hours at 4°C. Following ultracentrifugation, ∼500 µL fractions were collected from the bottom of the sucrose gradient tube using a Piston Gradient Fractionator (BioComp Instruments). Polysome profiles of each sample were assessed by continuous measuring of the absorbance at 260 (RNA) and 280 (proteins) nm using the Triax Flow Cell software (v1.56). After fractionation, TRIzol LS (Invitrogen) was added to each fraction (750 µL per 450 µL of sample), and the samples were stored at −80°C until RNA purification.

Area under the curve (AUC) was quantified for the pituitary polysome profiles in *R* (version 4.2.2) using *pracma* package to estimate ribosomal subunits and polysome-associated translational efficiency. Raw absorbance at 260 nm (Abs260) datasets were sorted in ascending order of fraction number and baseline correction was implemented. Specifically, the global minimum absorbance value across all fractions was defined as the baseline estimate. This value was subtracted from each Abs260 measurement to generate a baseline-corrected signal. Negative values resulting from this subtraction were truncated to zero to prevent artificial distortion of integrated areas. The polysome profiles (n = 2 replicates per condition, pool of 3 pituitaries in each replicate) were segmented into discrete translational regions corresponding to 40S, 60S, 80S (monosomes: 1 ribosome), and light (2-4 ribosomes) and heavy polysomes (>5 ribosomes). These regions were defined by determined fraction boundaries (xmin–xmax), reflecting characteristic peak distributions within sucrose gradients. AUC was calculated using the trapezoidal numerical integration method (via the *pracma::trapz* function), which approximates the integral of absorbance over fraction number. Finally, visualizations were generated using *ggplot2*.

### RNA Extraction

Fractions corresponding to monosomes, light polysomes, and heavy polysomes were pooled, and RNA was extracted from each group using TRIzol LS reagent (Invitrogen), following the phenol–guanidine isothiocyanate–chloroform extraction method. RNA was precipitated from the aqueous phase by adding isopropanol (0.5 mL per 0.75 mL of TRIzol LS), with 1 µL of glycogen (Thermo Fisher) added to each sample as a carrier to enhance RNA yield. Samples were incubated at **-**20°C overnight, then pelleted by centrifugation at 12000×g at 4°C for 15 minutes. The resulting RNA pellets were washed twice, first with 75% ethanol, then with 100% ethanol, and finally resuspended in 25 µL of nuclease-free water. RNA quantity and quality was measured by Nanodrop.

### RNA Sequencing

RNA was sent for high-throughput sequencing (Deep RNA-Seq) to Admera Health, USA. RNA quality was assessed by RNA Tapestation (Agilent Technologies Inc., California, USA) and quantified by Qubit 2.0 RNA HS assay (ThermoFisher, Massachusetts, USA). Poly (A) mRNA enrichment, library generation and sequencing was done by the company and as previously reported [1]. Briefly, paramagnetic beads coupled with oligo d(T)25 were combined with total RNA to isolate poly(A)+ transcripts based on NEBNext® Poly(A) mRNA Magnetic Isolation Module (New England BioLabs Inc., Massachusetts, USA). Prior to first strand synthesis, samples were randomly primed (5’ d(N6) 3’ [N=A,C,G,T]) and fragmented. The first strand was synthesized with the Protoscript II Reverse Transcriptase with a longer extension period, approximately 40 minutes at 42°C. All remaining steps for library construction were used according to the NEBNext® Ultra™ II Non-Directional RNA Library Prep Kit for Illumina® (New England BioLabs Inc., Massachusetts, USA). Final libraries quantity was assessed by Qubit 2.0 (ThermoFisher, Massachusetts, USA) and quality was assessed by TapeStation D1000 ScreenTape (Agilent Technologies Inc., California, USA). Final library size should be about 430 bp with an insert size of about 300 bp. Illumina® 8-nt dual-indices was used to increase the number of samples sequenced per run. Equimolar pooling of libraries was performed based on QC values and sequenced on an Illumina® Novaseq platform (Il-lumina, California, USA) with a read length configuration of 150 bp for 40M pair-end (PE) reads per sample (20M in each direction).

### Bioinformatics analyses of RNA-seq data

Quality of paired-end reads was assessed using FastQC and MultiQC. Reads were aligned to the *Mus musculus* reference genome (GRCm39, Ensembl latest release) using STAR (v2.7.11b). Resulting BAM files were sorted by genomic coordinates using the *--outSAMtype BAM SortedByCoordinate* option. Gene-level counts were generated from BAM files using featureCounts, with parameters set to include paired-end reads and primary alignments only.

### RNA-seq–based gene clustering, heatmap trajectory analysis, and GO enrichment across pituitary translatome fractions

To gain initial insight into genes representative of each translatome fraction, we identified transcripts with the highest relative expression across polysome fractions. Raw count matrices were imported into R (version 4.2.2) and processed using the DESeq2 package. Count data were rounded to integers and used to generate a DESeqDataSet with sample metadata specifying experimental variables, including genotype and polysome fraction. Variance stabilizing transformation (VST) was applied to normalized count data. Samples were grouped by fraction (input, monosome, light polysomes, and heavy polysomes), and VST-transformed expression values were averaged across biological replicates within each fraction.

Genes exhibiting the greatest variability across fractions were identified by variance ranking, and the top 2,000 most variable genes were selected for downstream analysis. To capture relative expression dynamics across fractions, gene-wise scaling (z-score normalization) was performed across the four fraction groups. Genes were subsequently clustered using k-means clustering (k = 6, nstart = 50) to identify distinct translational trajectories. Cluster assignments were used to generate gene trajectory heatmaps with the pheatmap package, where rows represented genes and columns represented polysome fractions. Cluster-level expression profiles were calculated by averaging scaled expression values within each cluster across fractions to characterize patterns of transcript distribution along the translational gradient.

Gene Ontology (GO) enrichment analysis was performed for each gene cluster to identify overrepresented biological processes. Gene lists corresponding to each cluster were filtered to retain valid gene symbols and analyzed using the clusterProfiler package with the Biological Process ontology. Statistical significance was assessed using Benjamini–Hochberg multiple testing correction, with a p-value cutoff of 0.05 and q-value cutoff of 0.2. Enriched GO terms were ranked based on adjusted p-values, and the top-ranked categories were retained for downstream visualization and interpretation. Cluster-specific biological programs were further annotated based on expression trajectory patterns across polysome fractions. All analyses were performed in R (version 4.4.2).

### PCA, heatmap, and differential expression analysis of transcriptome and translatome profiles in WT and *Magel2* KO pituitaries

Differential expression analysis was performed in R (version 4.2.2) using the DESeq2 package and raw RNA-seq count data. Comparisons between *Magel2* KO and WT samples were conducted independently for each fraction group (input, monosome, light polysome, and heavy polysome). Genes were considered differentially expressed at a threshold of p ≤ 0.0005 and an absolute log2 fold change (|log2FC|) ≥ 0.2. Gene identifiers were mapped to gene symbols using the org.Mm.eg.db annotation database through AnnotationDbi.

For heatmap visualization, a combined set of significant genes across all comparisons was selected using a threshold of p < 0.05 and |log2FC| ≥ 0.05. Variance stabilizing transformation (VST) was applied to normalized count data, and expression values were scaled by row (z-score normalization) to highlight relative expression differences across samples. Heatmaps were generated using the pheatmap package with sample annotation based on experimental condition.

Principal component analysis (PCA) was performed on VST-transformed expression data using the DESeq2 plotPCA function to assess sample variability and clustering. PCA plots were generated using ggplot2, with samples colored according to experimental condition. For downstream functional analyses, Gene Ontology (GO) enrichment analysis was performed using the enrichGO function in clusterProfiler, mapping ENSEMBL gene identifiers to Biological Process terms. GO categories with an adjusted p-value (padj) < 0.05 were considered significantly enriched. Gene Set Enrichment Analysis (GSEA) was subsequently performed using the same GO Biological Process ontology with a padj < 0.05 threshold. This analysis was further extended to Molecular Signatures Database (MSigDB) gene sets using the clusterProfiler package in R (version 4.2.2). When applicable, raw counts were normalized using the median-of-ratios normalization method in DESeq2 to generate normalized counts.

### Reverse transcription quantitative PCR (RT-qPCR) analysis

Gene expression levels were measured by RT-qPCR. A total of 100 ng of RNA isolated from input samples or pooled fractions corresponding to monosomes, light polysomes, and heavy polysomes was treated with DNase I (Roche, Basel, Switzerland or Qiagen, Hilden, Germany) to remove genomic DNA. RNA was then reverse-transcribed into cDNA using the High-Capacity cDNA Reverse Transcription Kit (Applied Biosystems, Waltham, MA, USA).

qPCR reactions were performed in technical triplicates using 10 ng of cDNA per well in 384-well plates on a QuantStudio 7 Pro Real-Time PCR System (Applied Biosystems) with PowerUp SYBR Green Master Mix (cat. #A25742, Applied Biosystems). Primers were designed and validated as previously described [57]. Primer sequences were as follows: mNdn-Q-F1, 5′-ACCTGAAGTACCAGCGTGTG-3′; and mNdn-Q-R1, 5′-ACGCCTGGGGATCTTTCTTG-3′. RT-qPCR data were analyzed using QuantStudio Design and Analysis 2 (DA2) software (Thermo Fisher Scientific) and Microsoft Excel. Baselines and threshold were set automatically to determine cycle threshold (Ct) values. Relative gene expression was calculated using the ΔCt method.

### Prolactin serum analysis

Serum was collected from 8–10-week-old WT and *Magel2* KO male mice under fed or 24-h fasting conditions (n = 6 per group; sacrifice at 1:00 P.M.). Serum prolactin levels were measured by the Endocrine Technologies Core at the Oregon National Primate Research Center using a commercial ELISA kit (Abcam). The assay detection range was 27.43–20,000 pg/mL, with intra- and inter-assay coefficients of variation of 12.5% and 19.6%, respectively.

### Plotting and statistical analyses

Unless indicated otherwise, GraphPad Prism Software (version 10.4.0) was used for all plotting and statistical analyses, and p-values ≤ 0.05 were considered statistically significant [*P* ≤ 0.05 (∗), *P* ≤ 0.01 (∗∗), *P* ≤ 0.001 (∗∗∗), *P* ≥ 0.05 (non-significant, ns)].

## RESULTS

### Proteomic analysis of *Magel2* KO pituitaries reveals importance of mRNA translation control

To gain initial insight into the role of Magel2 in the pituitary, we performed differential protein expression and enrichment analyses of previously published proteomic data from WT and *Magel2* KO pituitaries [39]. Downregulated proteins in *Magel2* KO pituitaries were enriched for translation- and ribosome-associated pathways, and co-translational SRP-dependent protein targeting, suggesting impaired translational regulation and defective targeting of secretory and membrane proteins (Figure 1A, B; Table S1). Based on these findings, we hypothesized that MAGEL2 contributes to translational regulation in the pituitary gland. We therefore aimed to optimize a polysome profiling workflow in mouse pituitaries and investigate the impact of *Magel2* loss on translation (Figure 1C).

**Figure 1.**
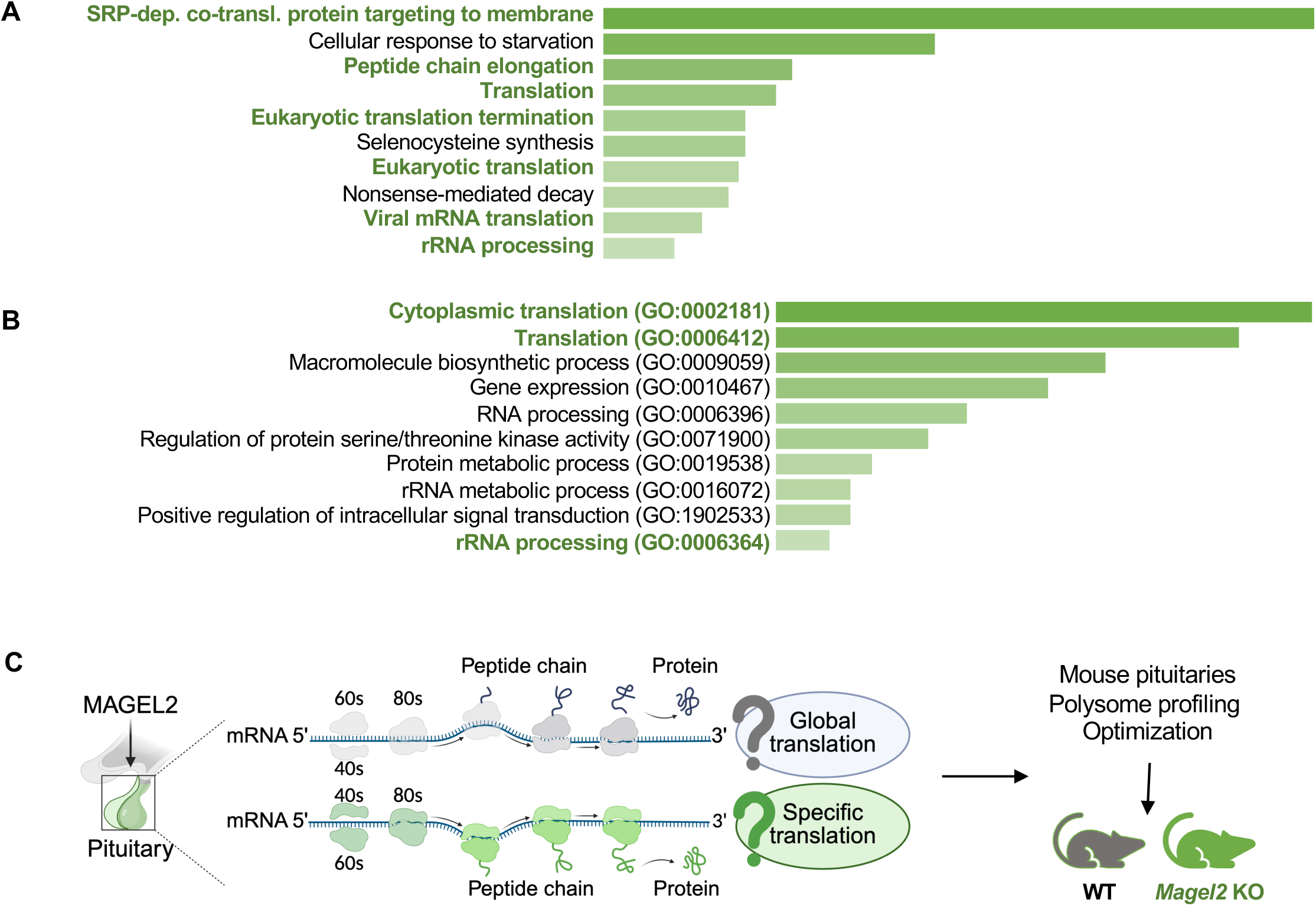
Gene set enrichment analysis of downregulated proteins in pituitaries of Magel2 knockout mice suggests impaired protein translation. Differential protein expression analysis was performed comparing wild type (WT) and Magel2 KO mice (n = 5 per group). Proteins with a log₂ fold change < −0.25 and p < 0.05 were selected for Enrichr analysis. Significantly enriched Reactome pathways **(A)** and GO Biological Process 2025 terms **(B)** are shown, ranked by p (see [38] and Table S1 for supporting data). **C**. Overview of the experimental design used to investigate the role of Magel2 in pituitary translation. The polysome profiling protocol was first optimized using varying numbers of pituitaries from WT mice, followed by profiling performed using three pituitaries per experiment.

### Polysome profiling optimization

Gene expression analyses in pituitaries of *Magel2* KO mice have been performed at the RNA and protein levels; however, the translational level and its regulation have not been explored so far [39, 58]. Polysome profiling is well established and a powerful technique to study the translational status of individual mRNAs by separation of polysomes, ribosomes, ribosome subunits (40S and 60S), and free (untranslated) mRNAs (Figure 2A) [1, 2, 48, 59]. Since there was no protocol available for running this experiment on the mouse pituitary, we first optimized the protocol for the mouse pituitary tissues based on a protocol that was previously described [2]. One of the major concerns in this study was the limited amount of starting material and determining how many individual glands to pool per sample.

**Figure 2.**
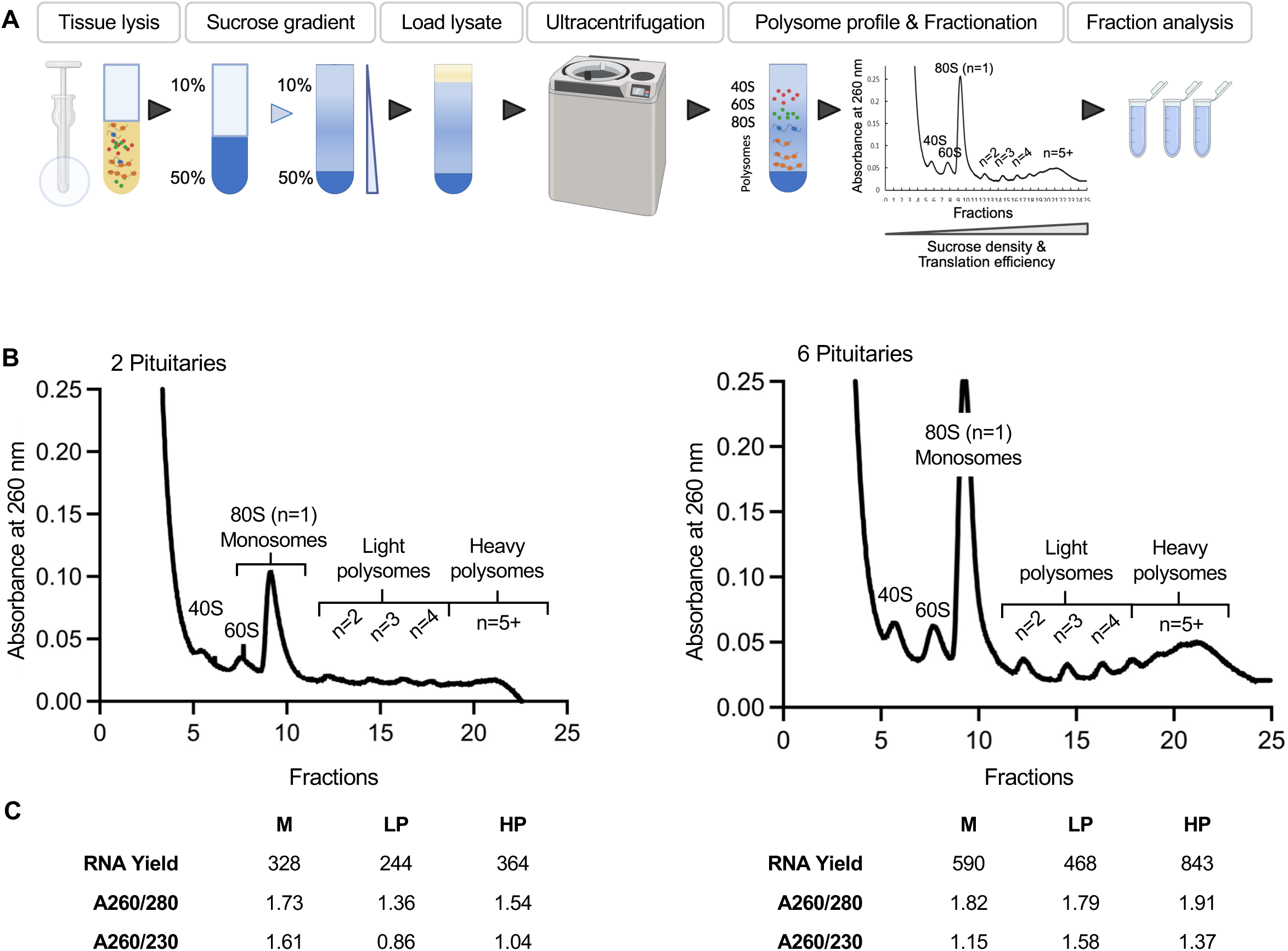
Experimental design for lysate preparation and polysome profiling. **A.** Overview of the method: tissue lysates are prepared and subjected to sucrose gradient ultracentrifugation to separate monosomes (single ribosomes) and polysomes (multiple ribosomes), followed by RNA extraction and downstream analysis of each fraction group [2]. **B.** Representative polysome profiles and **(C)** RNA yield (ng) obtained from each fraction group using 2 or 6 pituitaries.

To address this, we tested pools of 2 and 6 tissues for polysome profiling and RNA yield from fractions (Figures 2B and C). Our goal was to achieve clear polysome resolution and obtain sufficient, high-quality RNA from each fraction for downstream analyses. The polysome profiles from pituitary are demonstrated based on their A_260_ absorbance peaks based on whether RNA is associated with 40S, 60S, monosomes (80S - 1 ribosome), light polysomes (2-4 ribosomes), and heavy polysomes (and more than 5 ribosomes) (Figure 2B). We optimized polysome profiling using individual pituitaries as well as pooled samples from 2 and 6 pituitaries (Figure 2B). While sufficient polysome resolution was achieved in all conditions, RNA yield from some samples was borderline for downstream analyses. Considering profile quality, RNA yield, and the goal of minimizing animal use, we concluded that pooling 2-3 pituitaries yields sufficient material for reliable polysome profiling and downstream analysis.

### Polysome profiling of pituitary from WT and *Magel2* KO mice

After optimization, polysome profiling was performed on pituitaries from 8-week-old WT and *Magel2* KO male mice (n = 8 per group). Lysates were prepared from pooled tissues, with three animals combined for two technical replicates and two animals pooled for the third replicate. Polysome profiles were consistently well resolved across all samples (Figure 3A). Fractions were pooled into monosome, light polysome, and heavy polysome groups, and RNA isolated from each fraction, together with input RNA, was subjected to deep RNA sequencing (Figure 4A).

**Figure 3.**
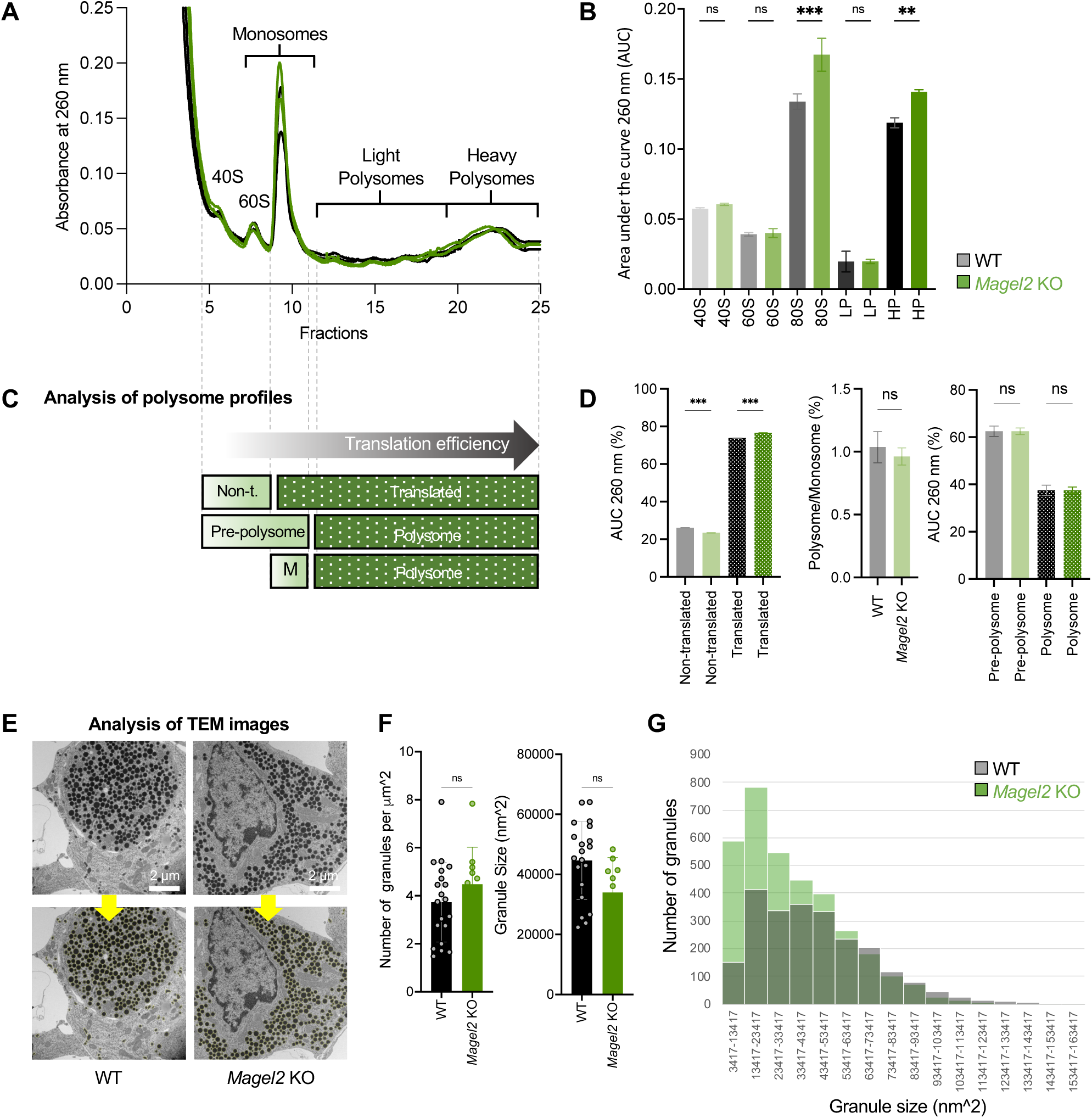
Analysis of the area under the curve in pituitary polysome profiles of Magel2 knockout mice suggests changes in translation efficiency. Polysome profiling was performed on pituitaries from WT and Magel2 KO mice (two biological replicates per group are shown, each consisting of a pool of three pituitaries). **A.** Representative pituitary polysome profiles for WT (black) and Magel2 KO (green), showing RNA distribution as absorbance at 260 nm (y-axis) across polysome fractions (x-axis). **B.** The area under the curve (AUC) at 260 nm for WT and *Magel2* KO profiles was calculated with baseline correction in R for the 40S, 60S, 80S (monosome), light polysome, and heavy polysome regions. **C-D.** Overview and quantification of the framework used to assess translation efficiency, including comparisons of non-translated (40S and 60S) versus translated (monosome, light, and heavy polysomes) regions; the polysome-to-monosome ratio; and pre-polysome (40S, 60S, and monosome) versus polysome (light and heavy polysomes) regions. **E-G.** Transmission electron microscopy (TEM) analysis of WT and Magel2 KO pituitary secretory granules (SGs) showing the total number, average size, and size distribution analysis of SGs in WT versus Magel2 KO pituitaries. Statistical significance was determined using the Brown–Forsythe and Welch ANOVA tests, followed by Dunnett’s T3 multiple comparisons test to compare WT and Magel2 KO groups. [P ≤ 0.01 (∗∗), P ≤ 0.001 (∗∗∗), P ≥ 0.05 (non-significant, ns)].

**Figure 4.**
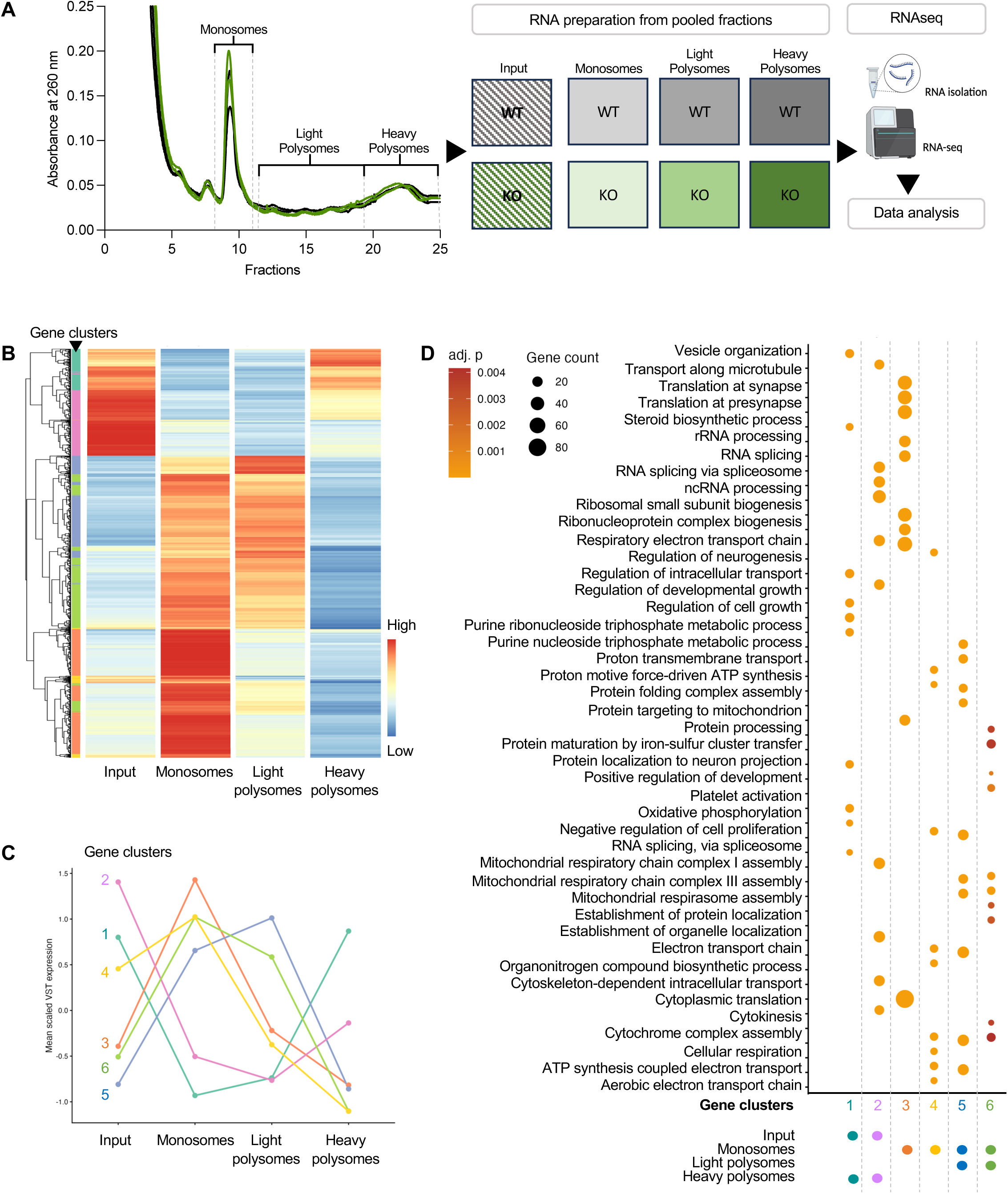
Polysome fractionation followed by RNA-seq reveals distinct gene clusters with differential translation patterns in the pituitary. **A.** Following polysome profiling of pituitaries from WT in gray and *Magel2* KO in green mice, fractions corresponding to similar profiles were pooled into three categories: monosome, light polysome, and heavy polysome for both genotypes. RNA was extracted from each pooled fraction, as well as from input samples, and subjected to RNA-seq and downstream analyses. **B.** Gene trajectory heatmap across input, monosome, light polysome, and heavy polysome fractions, showing six gene clusters based on variance stabilizing transformation (VST). **C.** Six gene clusters defined based on the similarity of VST values across the four fraction groups. **D.** Top Gene Ontology (GO) biological process terms associated with each cluster, including vesicle organization and regulation of developmental growth, translation at synapse, ribosome biogenesis, and mRNA splicing. See also Table S3.

To gain initial insight into the impact of *Magel2 l*oss on global translation in the pituitary, we first examined translatome changes by analyzing the area under the curve (AUC) of polysome profiles based on A260 absorbance peaks corresponding to the 40S, 60S, monosome, light polysome, and heavy polysome fractions (Figure 3B) [60]. We then calculated and compared sums of non-translated (40S and 60S) versus translated (monosome, light, and heavy polysomes) regions; the polysome-to-monosome ratio; and pre-polysome (40S, 60S, and monosome) versus polysome (light and heavy polysomes) regions (Figures 3C-D). Although the 80S and heavy polysome peak AUCs, as well as the comparison between non-translated and translated fractions, suggested a slight increase in *Magel2* KO pituitaries, comparisons of pre-polysome and polysome fractions and polysome-to-monosome ratios revealed no significant differences, indicating no major effect on global translation in *Magel2* KO pituitaries. Of note, these mice were maintained under non-stress conditions, warranting future studies under stress or fasting conditions.

Then we quantified SG number and size in WT and *Magel2* KO pituitaries (Figure 3E-G). SGs were identified on contrasted ultrathin TEM images as electron-dense, spherical structures with diameters ranging from 100–800 nm and clearly distinguishable boundaries from the surrounding cytoplasm [56]. Although total SG number and average granule size were not significantly different between groups, size distribution analysis suggested an increased proportion of smaller granules in *Magel2* KO pituitaries. The presence of more numerous smaller SGs suggests altered granule maturation and/or hormone trafficking dynamics in *Magel2* KO pituitary cells. Smaller secretory granules observed in the Magel2 KO pituitary may reflect impaired secretory capacity resulting from downregulation of SRP-dependent cotranslational protein targeting pathways [39]. Together with the polysome profiling results, these findings indicate that loss of Magel2 does not substantially affect overall translational capacity in the mouse pituitary but may influence translation and processing of specific transcripts.

### RNA-seq analysis of translatome fractions reveals distinct mRNAs enriched across different components of the translational machinery in the pituitary

To gain further insight into the pituitary translatome, we analyzed which mRNAs were associated with each fraction of the translational machinery (monosomes, light polysomes, and heavy polysomes) by performing RNA-seq on RNA isolated from the input and pooled fractions (Figure 4A). RNA-seq data were processed starting from raw FASTQ files (Figure S2A). After quality control, reads were mapped to the mouse reference genome using STAR, resulting in high-quality alignments with an average mapping rate above 80% (Figures S2B and S3). Then, we analyzed genes representative of the transcriptome (input) and each translatome fraction (monosome, light polysomes, and heavy polysomes) (Figure 4B-D) to identify transcripts with the highest relative expression across polysome fractions. Following normalization and variance stabilizing transformation (VST), the top 2,000 most variable genes across fractions were selected for downstream analysis (Figure 4B and ).

By applying gene-wise z-score normalization and k-means clustering, we identified 6 distinct gene clusters with translational trajectories across the polysome gradient (Figures 4B-C and Table S2 and S3). Cluster assignments revealed distinct transcript distribution trajectories across polysome fractions. Clusters 1 and 2 were enriched in the input and heavy polysome fractions, suggesting coordinated transcriptional and translational regulation, particularly of metabolic pathways (Figure 4D). In contrast, clusters 3 and 4 were enriched in monosomes, whereas clusters 5 and 6 were associated with monosome and light polysome fractions.

We then performed GO enrichment analysis and identified cluster-specific biological processes (Figure 4D and Table S3), with heavy polysome-enriched clusters showing pathways related to metabolism and oxidative phosphorylation. Interestingly, monosome-enriched clusters were strongly associated with RNA processing and translation-related processes (Figure 4D). This finding may reflect contributions from the posterior pituitary (neurohypophysis), which consists primarily of hypothalamic neuronal projections [61]. Monosome-associated translation has been shown to play an important role in neurons, where single ribosomes can locally synthesize proteins within spatially restricted neuronal and synaptic compartments [62]. Given that the posterior pituitary stores and releases hypothalamic peptide hormones, including oxytocin and vasopressin, these findings suggest that monosome-regulated translation may contribute to neuroendocrine function within the pituitary. Together, these data provide a resource for a better understanding of the mouse pituitary transcriptome and translatome, which will contribute to improved insight into translational regulation in neuroendocrine disorders, including PWS and SYS.

### Differential expression analysis of the transcriptome and translatome in WT and *Magel2* KO pituitaries

To evaluate transcriptomic and translatomic changes associated with *Magel2* depletion, we performed differential expression analysis (DEA) comparing WT and *Magel2* KO pituitary samples across input, monosome, light polysome, and heavy polysome fractions. First, to obtain an overview of gene expression across all samples in the four analyzed fraction groups, we generated a heatmap with hierarchical clustering based on gene expression, shown in Figure 5A. For heatmap generation and clustering, we selected genes with p ≤ 0.05 and an absolute log2 fold change ≥ 0.5 in comparisons between *Magel2* KO and WT within each fraction group. As shown in Figure 5A, input and heavy polysome samples clustered together, reflecting similar gene expression profiles, whereas monosome and light polysome samples formed a separate cluster. This pattern is consistent with our previous overall analysis of pituitary gene expression patterns shown in Figure 4B.

**Figure 5.**
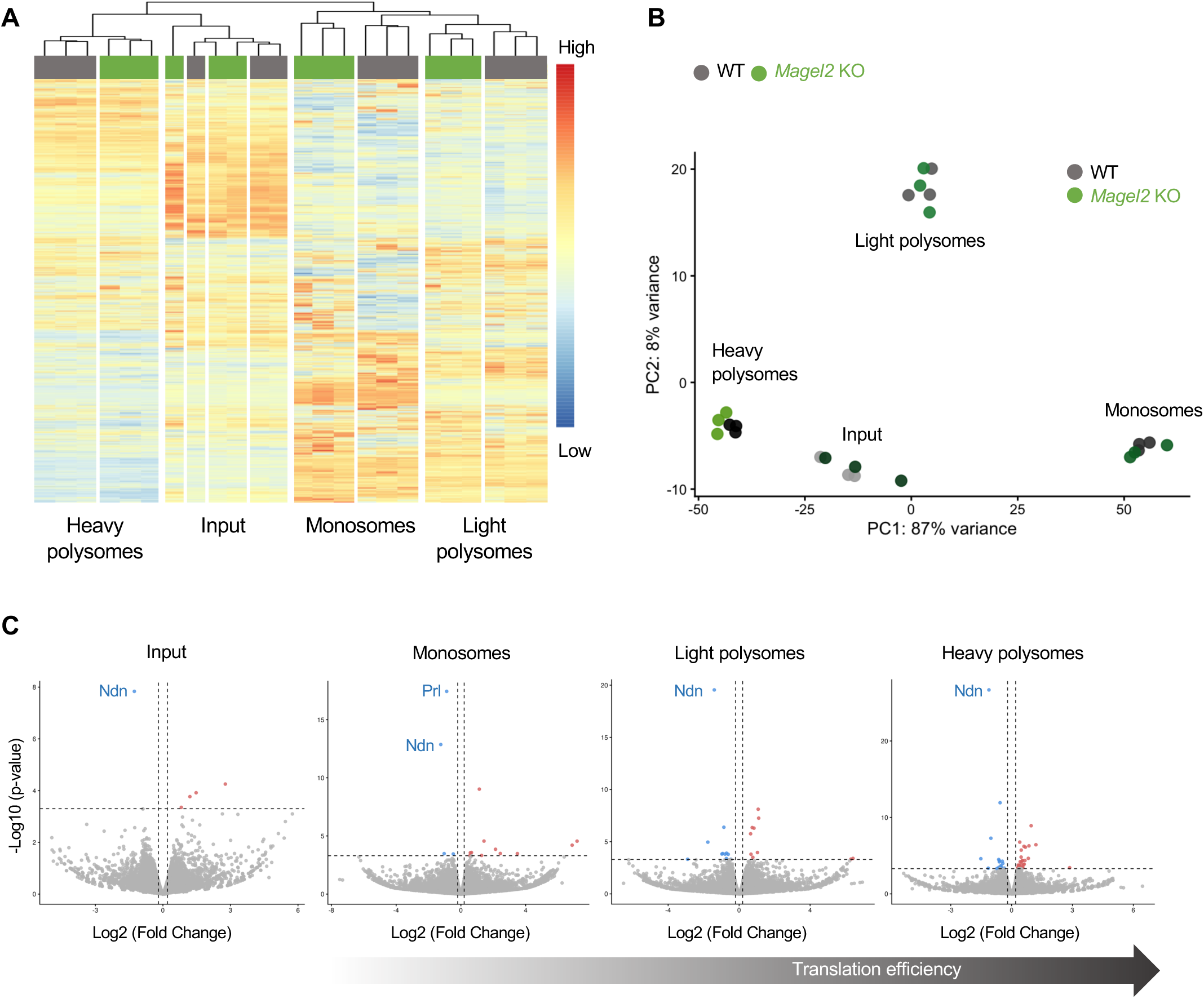
Differential expression analysis of transcriptome and translatome fractions in WT and Magel2 KO pituitaries. **A.** Heatmap of differentially expressed genes in the four fraction groups. Genes were selected based on p ≤ 0.05 and absolute log2 fold change ≥ 0.5 and ≤–0.5 between Magel2 KO in green and WT in gray within each fraction group. Expression values were scaled by row to visualize relative expression patterns. **B.** Principal component analysis (PCA) of samples. Each point represents a replicate, colored by experimental condition Magel2 KO in green and WT in gray. The first two principal components PC1 and PC2 are shown, with the percentage of variance explained indicated on each axis. **C.** Volcano plots showing genes that are significantly changed in the Magel2 KO mouse pituitary input, monosome, light, and heavy polysome fraction groups. The horizontal axes denote log2 fold change thresholds (≥ 0.2 and ≤–0.2) and vertical axes P value thresholds (P ≤ 0.0005).

We next performed principal component analysis (PCA) of samples based on variance stabilizing transformation (VST)-transformed expression data (Figure 5B). Input samples and each translatome fraction clustered into distinct groups containing both WT and *Magel2* KO samples. However, samples generally did not separate by genotype within most fraction groups, reflecting the relatively low number of differentially expressed genes in these fractions. In contrast, a slight separation between WT and *Magel2* KO samples was observed in the heavy polysome fraction, consistent with the higher number of differentially represented genes identified in this fraction, as shown by volcano plots (Figure 5C) and the bar graph (Figure 6A). Given the very limited number of differentially expressed genes (DEGs) reaching significant adjusted P-values, which is not unusual in biological samples, we applied less stringent thresholds of P ≤ 0.0005 and log2 fold change ≥ 0.2 or ≤ −0.2 for downstream analyses (Figures 5C and 6).

**Figure 6.**
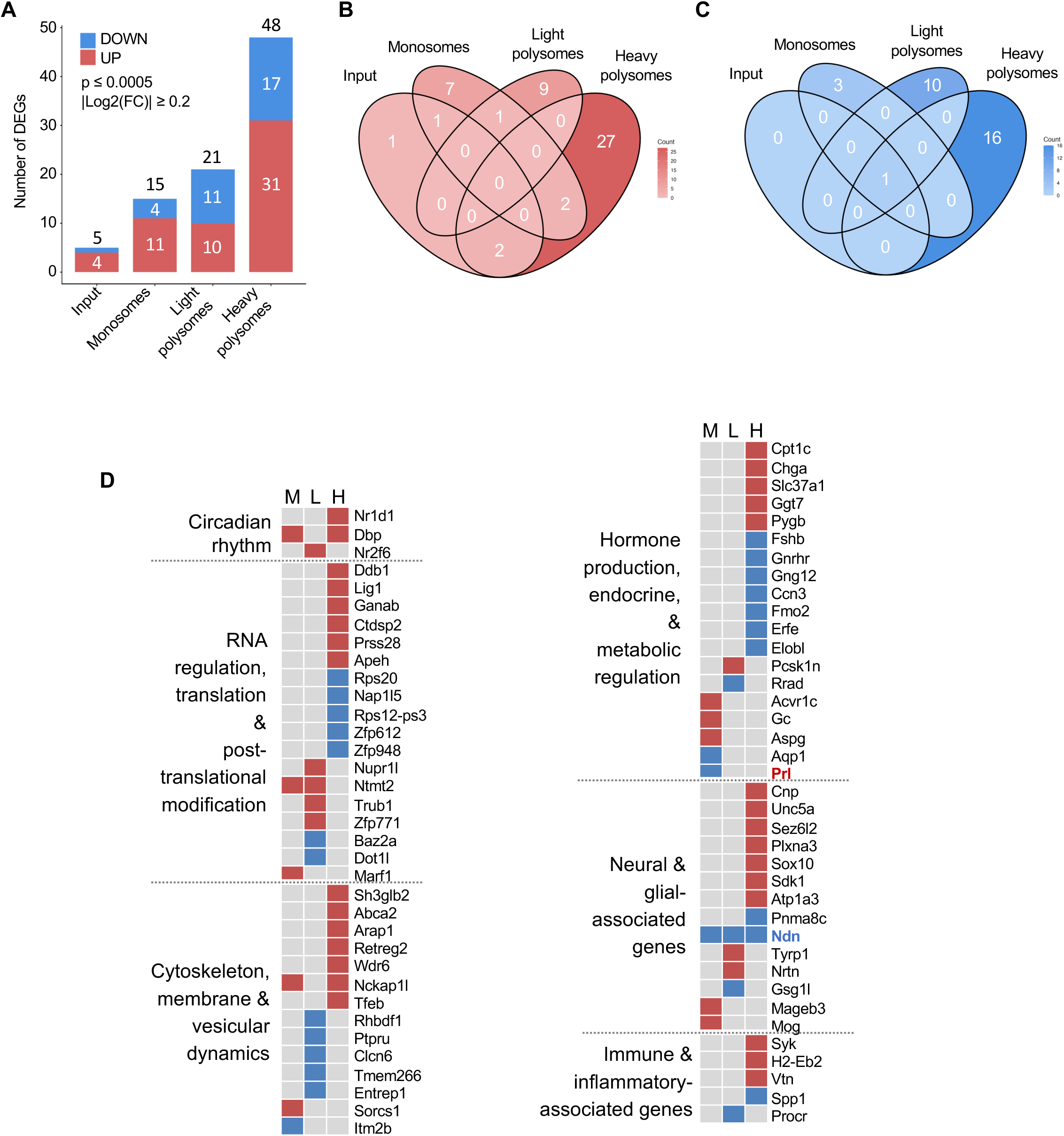
Differential expression analysis reveals enrichment of altered transcripts in the heavy polysome fraction of Magel2 KO pituitaries. **A.** Bar plots showing the number of significant downregulated in blue and upregulated in red transcripts in inputs, monosomes, light polysomes, and heavy polysomes with p ≤ 0.0005 and absolute log2 fold change ≥ 0.2 and ≤–0.2 cutoffs. **B-C.** Venn diagrams showing the downregulated blue and upregulated red differentially expressed and translated transcripts with p ≤ 0.0005 and absolute log2 fold change ≥ 0.2 across the four fraction groups. **D.** Functional annotation analysis of differentially translated transcripts identified downregulated (blue) and upregulated (red) genes with *p* ≤ 0.0005 and absolute log2 fold change ≥ 0.2 across monosome (M), light polysome (LP), and heavy polysome (HP) fractions. Enriched biological processes suggest that translational regulation has important functional relevance in the pituitary and may be altered in neuroendocrine disorders such as PWS and SYS.

To determine the overlap of differentially represented genes across fractions, we next analyzed their intersections between input and translatome fractions. Interestingly, *Necdin* (*Ndn*) was consistently downregulated across all fractions (Figures 5C and 6B-C). In contrast, most differentially represented genes were fraction-specific, with the strongest enrichment observed in the heavy polysome fraction (Figure 6B–C and Tables S2-3).

As depicted in Figure 6A, the number of DEGs increased across translatome fractions, whereas only a limited number of genes were altered at the transcriptional level between WT and *Magel2* KO pituitaries. DEGs identified in the input, monosome, light polysome, and heavy polysome fractions were subjected to Gene Ontology (GO) Biological Process enrichment analysis using clusterProfiler, as well as Canonical Pathway analysis using Ingenuity Pathway Analysis (IPA) (Figures S4 and S5). The identified pathways were subsequently organized into functional groups through expert-driven manual curation (Figure 6D and Table S4).

Based on these analyses, DEGs were categorized into groups reflecting key pituitary functions and processes associated with symptoms observed in PWS and SYS, including hormone production, endocrine and metabolic regulation, and circadian rhythm regulation [21, 63]. Interestingly, several altered genes were associated with cytoskeleton organization, trafficking, membrane organization, and vesicular dynamics, consistent with the previously described role of MAGEL2 in F-actin nucleation-mediated retrograde protein trafficking [42, 45] and regulated secretion of hormones [39]. In addition, multiple genes involved in RNA regulation, translation, and post-translational modification appeared to be regulated at the translational level, further supporting a role for MAGEL2 in translational regulation (Figure 6D).

To validate selected findings, we focused on some of the most interesting observations from this first translatome analysis of pituitaries from WT and *Magel2* KO mice, including the downregulation of the PWS-associated gene *Ndn* and alterations in prolactin translation (Figure 7). From the RNA-seq data, raw counts were first normalized, and RNA levels across distinct translatome fractions were plotted and compared with transcriptome (input) levels. We then analyzed gene distribution as a percentage of the total polysome fractions to gain better insight into potential translational regulation. Figures 7A and 7D show representative examples for Necdin and prolactin, respectively.

**Figure 7.**
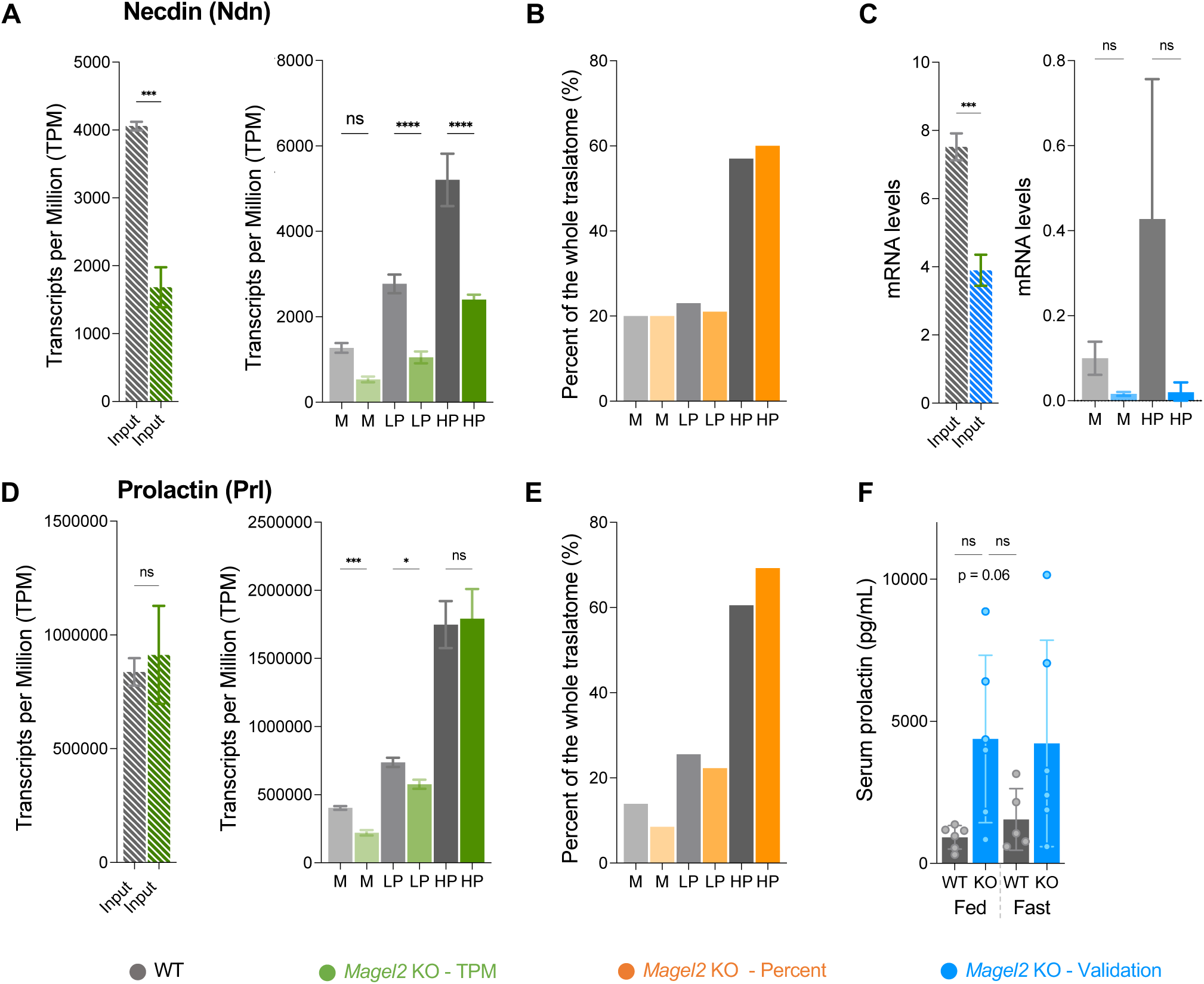
Validation of translatome data by RT-qPCR and proteome analyses. **A** and **D.** Normalized count values are shown for Necdin (Ndn) and Prolactin (Prl) in input and translatome fractions (M, monosome; LP, light polysome; and HP, heavy polysome fractions) of WT in gray and Magel2 KO in green (n = 3). **B** and **E.** Ratio of each fraction group (M, LP, and HP) in the whole translatome (M+LP+HP) is shown for Ndn and Prl in WT in gray and Magel2 KO in orange. **C.** RT-qPCR analysis of Ndn mRNA levels in the input and translatome fractions in WT in gray and Magel2 KO in blue. Data are presented as 2^ΔCt values with mean ± standard deviation from three biological replicates for polysome samples or three technical replicates for the inputs. **F.** Serum analysis of prolactin in WT in gray and Magel2 KO in blue in 8–10-week-old male mice under fed and 24-hour fasting (fast) conditions (n = 6 per group). Statistical significance was determined using a t-test for input samples or Brown–Forsythe and Welch ANOVA followed by Dunnett’s T3 multiple comparisons test for WT and Magel2 KO groups. Significance levels were defined as P ≤ 0.01 (∗∗), P ≤ 0.001 (∗∗∗), and P ≥ 0.05 (non-significant, ns).

These findings were further validated by RT-qPCR and ELISA analyses (Figures 7C and 7F). Together, our data demonstrated that *Ndn* is transcriptionally downregulated in pituitaries of *Magel2* KO mice, resulting in significantly lower levels of translation-associated transcripts (Figures 7A-C). In contrast, prolactin transcript levels were unchanged in the input fraction; however, decreased representation in monosome and light polysome fractions suggested increased engagement with heavy polysomes and enhanced translation. This observation was consistent with the increased serum prolactin levels detected in *Magel2* KO mice (Figures 7D-F). Alternatively, decreased engagement with monosomes may also reflect impaired translation, potentially due to reduced levels of SRP components, as observed in our previous proteomics analysis [39]. This could be associated with lower prolactin protein levels in the pituitary, although it remains unclear whether the affected forms represent mature or immature Prl protein. Together, these findings suggest that Magel2 impacts pituitary function at both the transcriptional and translational levels. Further studies are warranted to identify the molecular mechanisms underlying these changes and to determine their physiological relevance and contribution to PWS and SYS under stress conditions, such as fasting and metabolic stress.

## DISCUSSION

Our study provides the first comprehensive characterization of the mouse pituitary translatome and suggests that Magel2 translation of selected mRNAs in the mouse pituitary gland. Although translational regulation is increasingly recognized as a critical mechanism controlling endocrine activity enabling rapid hormonal responses without requiring new mRNA synthesis, its contribution to pituitary physiology has remained largely unexplored. By combining polysome profiling with RNA sequencing and TEM analysis, we demonstrated that loss of *Magel2* induces substantial alterations in polysome-associated RNAs despite relatively limited transcriptomic changes, suggesting that translational regulation constitutes an important layer of gene expression control downstream of MAGEL2 function.

The pituitary serves as the endocrine core of the organism, regulating hormone synthesis and secretion to adapt to changing metabolic and reproductive demands. Hypothalamic–pituitary neuroendocrine dysfunction is a hallmark of PWS and SYS and contributes to phenotypes including hyperphagia, hypogonadism, growth retardation, and hypothyroidism [21]. Given that *MAGEL2* is most highly expressed in the hypothalamus [21], previous studies have primarily linked MAGEL2 protein to hypothalamic neuroendocrine regulation, while pituitary dysfunction has largely been considered secondary to hypothalamic defects and therefore has not been investigated in detail. MAGEL2 deficiency has been associated with altered endosomal protein recycling and impaired neuropeptide maturation and secretion, however, whether MAGEL2 directly influences translational processes has remained unclear [39, 42, 45]. Our findings extend the functional significance of MAGEL2 beyond its established role in protein trafficking, secretory granule formation, and secretion, and suggest that MAGEL2 may also play an important role in regulating protein synthesis. Furthermore, our data also suggest that MAGEL2 deficiency in the pituitary may contribute directly to the endocrine phenotypes observed in PWS and SYS.

PWS is caused by chromosomal deletions, imprinting defects, or uniparental disomy of chromosome 15q11.2–q13, a region encompassing several imprinted protein-coding genes and noncoding RNAs, including *MAGEL2* [21]. Pathogenic mutations in *MAGEL2* cause SYS, which shares several clinical features with PWS [63]. Importantly, patients carrying deletions affecting only *MAGEL2* can still exhibit core PWS phenotypes, suggesting a critical contribution of MAGEL2 to symptom development in both disorders and raising important questions regarding the molecular mechanisms underlying MAGEL2 function [64]. Over the last decade, several molecular and cellular aspects of MAGEL2 molecular function have been uncovered. MAGEL2 is a member of melanoma antigen gene family (MAGE) which were shown to bind to E3 ubiquitin ligases and regulate ubiquitination of target proteins [40, 57]. Together with the E3 ubiquitin ligase TRIM27 and USP7 (MUST complex) [40], MAGEL2 ubiquitinates the WASH complex, through which MAGEL2 regulates protein localization [42, 45] and neuropeptide and hormone secretion in the hypothalamus [39]. MAGEL2 was also found to regulate the ubiquitination, stability, and nuclear-cytoplasmic distribution of CRY1 through interactions with RBX1 and USP7 [65]. Recently, MAGEL2 was found to shuttle between the nucleus and cytoplasm, and its N-terminal region may regulate m6A-modified mRNA metabolism [23], suggesting additional functions of MAGEL2. Here, we investigated the role of MAGEL2 in protein translation in the mouse pituitary. This work was motivated by our previous proteomics data indicating altered protein translation in *Magel2* KO mice, a model that recapitulates features of PWS and SYS [39]. Furthermore, dysregulated translational and ribosomal processes have also been linked to PWS neurons, suggesting altered ribosome- and translation-related pathways in PWS [22, 66]. We optimized polysome profiling, a well-established and widely used technique to study protein translation based on separation of RNA on sucrose gradients and its association with ribosomes. Although this method has been applied to multiple tissues, including the brain, to our knowledge it has not previously been optimized for the pituitary [2]. This study thus represents the first analysis of translatome in the mouse pituitary and alterations upon *Magel2* deletion, what is relevant to PWS and SYS.

First, we evaluated global polysome profiles by analyzing the area under the curve (AUC) of A260 absorbance peaks corresponding to the 40S, 60S, monosome, light polysome, and heavy polysome fractions (Figure 3A-D). The results did not reveal major differences in overall translational activity between WT and *Magel2* KO pituitaries, suggesting that loss of Magel2 does not induce a global translational shutdown or broad impairment of ribosome loading. Instead, the observed effects likely reflect selective translational regulation of specific transcripts and pathways. Such transcript-specific translational control is increasingly recognized as a mechanism that enables rapid endocrine adaptation while maintaining global protein synthesis homeostasis [67].

Given our previous findings that Magel2 affects SG number and size in the hypothalamus [38], we next examined SG ultrastructure in pituitary cells. Although total SG number and average granule size were not significantly different between groups, size distribution analysis suggested an increased proportion of smaller granules in *Magel2* KO pituitaries (Figure 3E-G). The presence of more numerous but smaller SGs may reflect several, not mutually exclusive, biological changes related to hormone synthesis, packaging, maturation, trafficking, or secretion in *Magel2* KO pituitary cells [68]. Together with the polysome profiling results, these findings suggest that loss of Magel2 does not substantially impair overall translational capacity in the pituitary, but rather may selectively influence the translation, processing, trafficking, and secretory handling of specific transcripts and their protein products.

Consistent with this interpretation, fraction-specific translatome analyses identified distinct translational trajectories across monosome and polysome fractions, indicating that pituitary RNAs are differentially partitioned according to their translational status (Figure 4). Heavy polysome-associated transcripts were enriched for metabolic and oxidative phosphorylation pathways, whereas monosome-associated transcripts were linked to RNA processing and translation-related functions. These findings suggest the existence of specialized translational programs within pituitary cells and support the concept that translational regulation contributes to endocrine functional specialization, allowing for fast adaptation to the needs and scaling up of the protein synthesis [69].

Notably, translatomic alterations in *Magel2* KO pituitaries were substantially more pronounced than transcriptomic changes, particularly within heavy polysome fractions. This suggests that translational regulation may represent an important mechanism through which MAGEL2 influences pituitary physiology. Functional enrichment analyses identified dysregulation of pathways associated with key pituitary physiological functions impaired in PWS and SYS. Many of these processes are closely interconnected with hormone synthesis and secretion and include pathways previously reported to be altered in patients and mouse models of PWS and SYS [21, 25], such as endocrine signaling and circadian rhythm regulation. Furthermore, we observed alterations in cellular processes closely linked to known MAGEL2 functions, including cytoskeleton organization, vesicular trafficking, secretory granule formation [39, 42, 45]. Intriguingly, several transcripts involved in protein translation were altered, further supporting the relevance of translational regulation in MAGEL2-associated pathologies.

For example, *Gnrhr* and *Fshb* were downregulated in the heavy polysome fractions of *Magel2* KO pituitaries (Figure 6), suggesting impaired gonadotroph function and dysregulation of the hypothalamic-pituitary-gonadal axis that may contribute to hypogonadism. *Gnrhr* encodes the gonadotropin-releasing hormone receptor that mediates hypothalamic GnRH signaling in pituitary gonadotrophs, whereas *Fshb* encodes the beta subunit of follicle-stimulating hormone (FSH). Although altered FSH levels have been reported in PWS, the mechanisms regulating FSH synthesis and secretion remain poorly understood [70–72].

Another particularly interesting finding was the altered monosome and light polysome association of prolactin (*Prl*) transcripts in the absence of significant transcript-level changes, suggesting altered translational regulation of *Prl* mRNA (Figure 7D–F). PRL is a secretory protein containing an N-terminal signal sequence recognized by signal recognition particles (SRPs). Association of the nascent signal peptide with SRP is critical for efficient translation and proper targeting to the endoplasmic reticulum. Failure of this interaction can trigger mRNA degradation through the recently described regulation of aberrant protein production (RAPP) pathway [44]. Although prolactin protein levels were reduced in pituitary proteomic analyses from the same mouse model [39], we found that circulating serum prolactin levels were elevated in *Magel2* KO mice. This finding may suggest increased prolactin synthesis and secretion and is consistent with reports of hyperprolactinemia in a subset of patients with PWS [73]. Alternatively, elevated circulating prolactin together with reduced pituitary prolactin levels may reflect increased hormone release and depletion of pituitary prolactin stores [73]. Together, these findings suggest that MAGEL2 may play multiple roles in regulating prolactin synthesis and/or secretion in the pituitary and warrant further investigation into the underlying molecular mechanisms. Although routine prolactin measurements have been recommended in PWS, the mechanisms underlying prolactin alterations remain poorly understood.

We also identified dysregulation of circadian rhythm–related transcription factors in the *Magel2* KO pituitary translatome, including the nuclear receptors *Rev-Erbα* (*Nr1d1*) and *Nr2f6,* as well as the transcription factor *Dbp,* suggesting that Magel2 contributes to regulation of the core circadian clock within the pituitary. Although MAGEL2 is well established as a circadian gene required for fine-tuning daytime activity, and in cells and neurons MAGEL2 has been shown to regulate BMAL1, PER2, and CRY1 [45, 54, 82, 83], its role in pituitary circadian regulation has remained unexplored. Supporting this connection, Nr1d1 loses its normal day/night differential expression in the hypothalamus of *Ndn–Magel2* double knockout mice [58]. Together, our findings suggest that MAGEL2-dependent translational regulation contributes to neuroendocrine and circadian rhythm dysfunction in PWS/SYS.

Interestingly, we identified several components of the translational machinery dysregulated in the *Magel2* KO pituitary translatome (Figure 6), further supporting a role for MAGEL2 in translational regulation. The transcription factor Creb3l2 was identified as a scaling factor for translation capacity in pituitary secretory cells during development, regulating approximately 75% of translation-related genes [74]. However, little is known about specialized translational regulation in the adult pituitary. Our data suggest that MAGEL2 may contribute to this process. Interestingly, our data show that Tfeb is upregulated in the heavy polysome fractions of the *Magel2* KO pituitary. Although TFEB is best known as a master transcriptional regulator of lysosomal biogenesis and autophagy, increasing evidence suggests that it also influences translational capacity and ribosome function [75]. Together, these findings suggest that MAGEL2 contributes to reprogramming of the translational machinery in the pituitary, potentially regulating the translation of specific mRNAs at both the transcriptional and translational levels through mechanisms involving ribosomal heterogeneity and specialized ribosome function.

Lastly, it was notable that Gene Ontology (GO) terms enriched in the differential translatome included protein trafficking and vesicle transport, processes in which MAGEL2 function is well established. For example, levels of Nck-associated protein 1–like (*Nckap1l*) were increased. Nckap1l is an essential component of the Wiskott–Aldrich syndrome protein family verprolin-homologous protein (WAVE) complex, whose members are key activators of the actin-related protein (ARP) 2/3 complex [76]. This function is analogous to that of the WASH complex, a known ubiquitination target of MAGEL2 [42]. These findings suggest the existence of a compensatory mechanism that promotes actin polymerization in the absence of Magel2.

In parallel, we analyzed the pituitary transcriptome of WT and *Magel2* KO mice as input for the translatome analysis and made the intriguing discovery that another PWS-associated gene, *Necdin* (*Ndn*), was significantly downregulated in the pituitaries of *Magel2* KO mice. *NDN* and *MAGEL2* belong to the type II *MAGE* gene family and are highly expressed in the pituitary and hypothalamus [21, 57]. *Ndn* has previously been detected in the developing mouse pituitary and proposed to play a role in pituitary development [87, 88]. This finding was unexpected, as *Ndn* has been proposed to compensate for *Magel2* loss because of their shared membership in the MAGE protein family. Indeed, *Ndn* is upregulated in the hypothalamus of *Magel2* KO mice, while conversely, increased *Magel2* expression has been reported in the hypothalamus of *Ndn* KO mice [58], suggesting reciprocal regulation and coordinated expression of these two *MAGE* genes in the hypothalamus. In contrast, we found that *Ndn* was downregulated in the *Magel2* KO pituitary at both the transcriptome (input) and translatome levels (Figure 7A), indicating that its regulation likely occurs at the transcriptional level. The consistent reduction of *Ndn* expression in the absence of *Magel2* suggests that these genes may also be co-regulated in the pituitary and raises the possibility that *MAGEL2* influences the expression of additional genes within the PWS locus and that this regulation is tissue specific.

Overall, our findings suggest that MAGEL2 impacts pituitary physiology, likely not only indirectly through hypothalamic regulation but also through direct effects within the pituitary itself. Furthermore, our data imply that MAGEL2 has multiple cellular functions at several levels, extending beyond its previously established role in K63-linked ubiquitin-mediated protein homeostasis to include newly identified functions in transcriptional and translational regulation. This is consistent with a recent multi-omics analysis of cortical neurons derived from CRISPR/Cas9-engineered hiPSC lines modeling SYS and PWS, which suggested that although a subset of dysregulated proteins is directly associated with MAGEL2-related ubiquitination, a substantial proportion of SYS/PWS-associated phenotypes may arise from broader proteome-level alterations [77], supporting the existence of additional MAGEL2 functions. Our work further expands this concept by identifying MAGEL2-dependent translational regulation in the pituitary, opening new avenues for uncovering previously unrecognized cellular and molecular functions of MAGEL2 that may ultimately reveal novel therapeutic targets.

## STUDY LIMITATIONS

In this study, we examined differentially translated genes associated with monosome, light polysome, and heavy polysome fractions. While polysome profiling is a powerful tool for assessing the translational state at a given time point, we acknowledge that it does not directly measure active translation or capture the temporal dynamics of polysome assembly and disassembly. Addressing these questions will require additional approaches such as ribosome profiling or puromycin-based assays and metabolic labeling.

Furthermore, the molecular mechanisms by which MAGEL2 impacts translation, including potential effects on ribosomal composition or direct association with ribosomes, remain to be determined. Because most experiments were performed in male mice, the results may not fully capture sex-specific differences in pituitary translation arising from the sexually dimorphic nature of the gland. Future studies including female mice will therefore be important to determine whether translational profiles differ between sexes. Given that mouse models of PWS and SYS exhibit abnormal feeding behavior, endocrine dysfunction, and metabolic dysregulation, and considering the established role of MAGE proteins in stress adaptation, further investigation of translational regulation under nutrient deprivation and metabolic stress conditions is warranted. Lastly, comparative studies across species will be important to better understand species-specific expression and functions of MAGEL2 within the hypothalamic–pituitary axis [57].

## Author Contributions

Conceptualization, T.B., Z.K. and K.F.T.; methodology, T.B., E.B.T., C.C.R.A., Z.K., M.C.H.S., F.Y.P., S.M., M.V., and K.F.T.; validation, T.B., F.Y.P.; data analysis, T.B., D.Š., J.S.S.G., C.C.R.A., M.C.H.S., E.B.T.; investigation, T.B.; resources, Z.K., A.L.K., E.B.T., K.F.T.; data curation, T.B., E.B.T., J.S.S.G., C.C.R.A., M.C.H.S., D.Š., Z.K., and K.F.T.; writing—original draft preparation, T.B., and K.F.T.; writing—review and editing, T.B., and K.F.T.; visualization, T.B., D.Š., and K.F.T.; supervision, K.F.T. and Z.K.; project administration, K.F.T. and Z.K.; funding acquisition, K.F.T. and Z.K. All authors have read and agreed to the published version of the manuscript.

## Institutional Review Board Statement

### Data Availability Statement

The raw and processed data supporting the conclusions of this article are available in the Gene Expression Omnibus database from the NCBI under accession number GS GSE330453

## Acknowledgments

We thank members of the Fon Tacer and Rosa laboratories for their advice and critical discussions. TEM sectioning and imaging were performed by Stephanny Lizarraga at the Texas Tech University College of Arts & Sciences Microscopy Center (CASM). Part of figures 1c and 2a were created in BioRender.com.

