## Supplementary material for "Loss of MAGEL2 Disrupts Pituitary Translation in a Mouse Model of PWS and Schaaf–Yang Syndrome": Sup Methods

**Polysome Profiling Reveals the Pituitary Translatome and Uncovers the Role of MAGEL2 in mRNA Translation in a Prader–Willi/Schaaf–Yang Mouse Model**

Tara Bayat^1,2^, Maria Camila Hoyos Sanchez^1,2^, Cristian Camilo Rodríguez-Almonacid^3^, Elena B Tikhonova^3^, Juan Sebastian Solano Gutierrez^1,2,4^, Denis Stepihar^1,2^, Farzana Yeasmin Popy^1,2^, Stephanie Myers^1,2^, Miloš Vittori^5^, Andrey L. Karamyshev^2,3^, Zemfira N. Karamysheva^2,3*^, and Klementina Fon Tacer^1,2*^

^1^School of Veterinary Medicine, Texas Tech University, Amarillo, TX 79106, USA

^2^Texas Center for Comparative Cancer Research (TC3R), Amarillo, TX 79106, USA
^3^Department of Cell Biology and Biochemistry, Texas Tech University Health Sciences Center, Lubbock, TX 79430, USA

^4^Center for Biotechnology and Genomic Medicine, Medical College of Georgia, Augusta University, Augusta, GA 30912, USA

^5^ University of Ljubljana, Biotechnical Faculty, Department of Biology, Ljubljana, Slovenia

**SUPPLEMENTARY FIGURE LEGENDS**

**Figure S1. Overview of Magel2 KO mouse breeding strategy, genotyping, and body and tissue weights.**

**A.** Breeding strategy used to generate experimental cohorts. Due to maternal imprinting and silencing of the maternal allele, offspring inheriting a paternal deletion of Magel2 (Magel2^pΔ/m+^) are functional knockouts, whereas mice carrying a maternal deletion are phenotypically WT. **B.** Genotyping of mice by PCR, showing representative detection of WT and Magel2 KO alleles. **C.** Body and tissue weights of animals used in the study.

**Figure S2. Data analysis pipeline of the RNAseq analysis of the polysome profiling practions.**

**A.** Schematic overview of the analysis pipeline. **B.** Quality control summary of pituitary RNA-seq reads based on FastQC analysis. Per-base sequence quality scores across all read positions show consistently high mean Phred scores across the entire read length, predominantly above Q30, indicating high-quality sequencing data suitable for downstream transcriptomic and translatomic analyses.

**Figure S3. Mapping and counting summary of pituitary RNA-seq reads.**

**A.** Mapping summary showing the proportion of unmapped (dark gray), multi-mapped (light gray), and uniquely mapped reads (pink) across all samples. **B.** Counting summary showing the proportion of reads assigned (pink) or unassigned (gray) to annotated gene features following featureCounts analysis. The majority of reads were successfully assigned to genes, supporting the quality and reliability of downstream differential expression and translatome analyses.

**Figure S4. GO Biological Process enrichment of differentially expressed genes across fractions.**

**A.** Gene Ontology (GO) Biological Process enrichment analysis was performed on significantly upregulated (up) and downregulated (down) genes identified in input, monosome, light polysome, and heavy polysome fractions. Differentially expressed genes were defined using thresholds of |log₂ fold change| ≥ 0.2 and p-value ≤ 0.0005. Enrichment analysis was conducted using *clusterProfiler* with the mouse reference database (*org.Mm.eg.db*), using dataset-specific gene sets as background. The plot displays a curated set of biologically relevant GO terms derived from the enrichment results. Each dot represents a significantly enriched pathway, with dot size proportional to the number of genes contributing to the enrichment (gene count). Pathways are separated by regulation direction (upregulation and downregulation), and colored accordingly (red for upregulation, blue for downregulation). The x-axis indicates the dataset (input, monosome, light polysome, heavy polysome), and the y-axis shows the selected GO Biological Process terms. **B.** Heatmap showing the top Canonical pathways identified by Ingenuity Pathway Analysis across all four groups. Pathways were included if they met the significance criteria (p-value < 0.05, |log2FC| > 0.2) in at least one group. Each row corresponds to a Canonical pathway, and each column corresponds to a group. Color intensity reflects the activation z-score, with blue indicating inhibition and red indicating activation.

**Figure S5. GO Biological Process enrichment of differentially expressed genes across fractions.**

Gene Ontology (GO) Biological Process enrichment analysis was performed on significantly upregulated (up) and downregulated (down) genes identified in input, monosome, light polysome, and heavy polysome fractions. Differentially expressed genes were defined using thresholds of |log₂ fold change| ≥ 0.2 and p-value ≤ 0.0005. Enrichment analysis was conducted using *clusterProfiler* with the mouse reference database (*org.Mm.eg.db*), using dataset-specific gene sets as background. The plot displays a curated set of biologically relevant GO terms derived from the enrichment results. Each dot represents a significantly enriched pathway, with dot size proportional to the number of genes contributing to the enrichment (gene count). Pathways are separated by regulation direction (upregulation and downregulation), and colored accordingly (red for upregulation, blue for downregulation). The x-axis indicates the dataset (input, monosome, light polysome, heavy polysome), and the y-axis shows the selected GO Biological Process terms.

**Supplementary Tables**

**Table S1. Gene set enrichment analysis of downregulated proteins in pituitaries of Magel2 KO mice suggests impaired protein translation.**

Differential protein expression analysis was performed comparing wild-type (WT) and Magel2 KO pituitaries (n = 5 per group). Proteins with a log₂ fold change < −0.25 and p < 0.05 were selected for Enrichr analysis. Significantly enriched Reactome pathways (A) and GO Biological Process 2025 terms (B) are shown, ranked by p value (see Figure 1A and B).

**Table S2 and S3. Clustering and functional enrichment analysis of transcriptome and translatome profiles across polysome fractions.**

RNA-seq data from the transcriptome (input) and translatome fractions (monosome, light polysomes, and heavy polysomes) were analyzed to identify transcripts with the highest relative expression across polysome fractions. Following normalization and variance stabilizing transformation (VST), the top 2,000 most variable genes were selected for downstream analyses. Gene-wise z-score normalization and k-means clustering identified six distinct gene clusters with characteristic translational trajectories across the polysome gradient (Table S2). GO enrichment analysis identified cluster-specific biological processes (Table S3).

**Table S4. Gene Ontology (GO) Biological Process enrichment analysis of differentially expressed genes across transcriptome and translatome fractions.**

GO Biological Process enrichment analysis was performed using clusterProfiler for differentially expressed genes (DEGs) identified in the input, monosome, light polysome, and heavy polysome fractions. Canonical pathway analysis was additionally performed using Ingenuity Pathway Analysis (IPA).
