## Supplementary figures and images for "Loss of MAGEL2 Disrupts Pituitary Translation in a Mouse Model of PWS and Schaaf–Yang Syndrome"

### Sup Figures

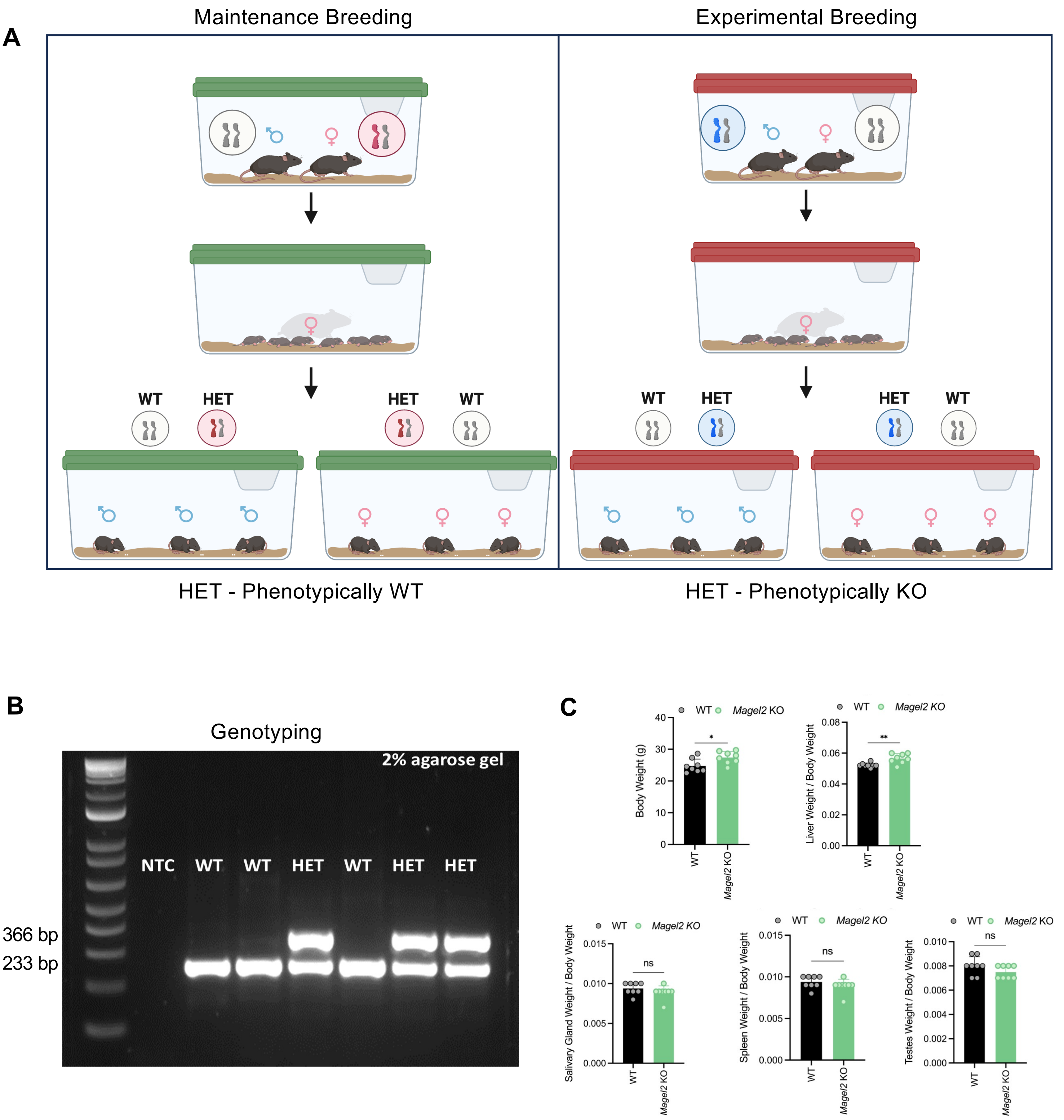

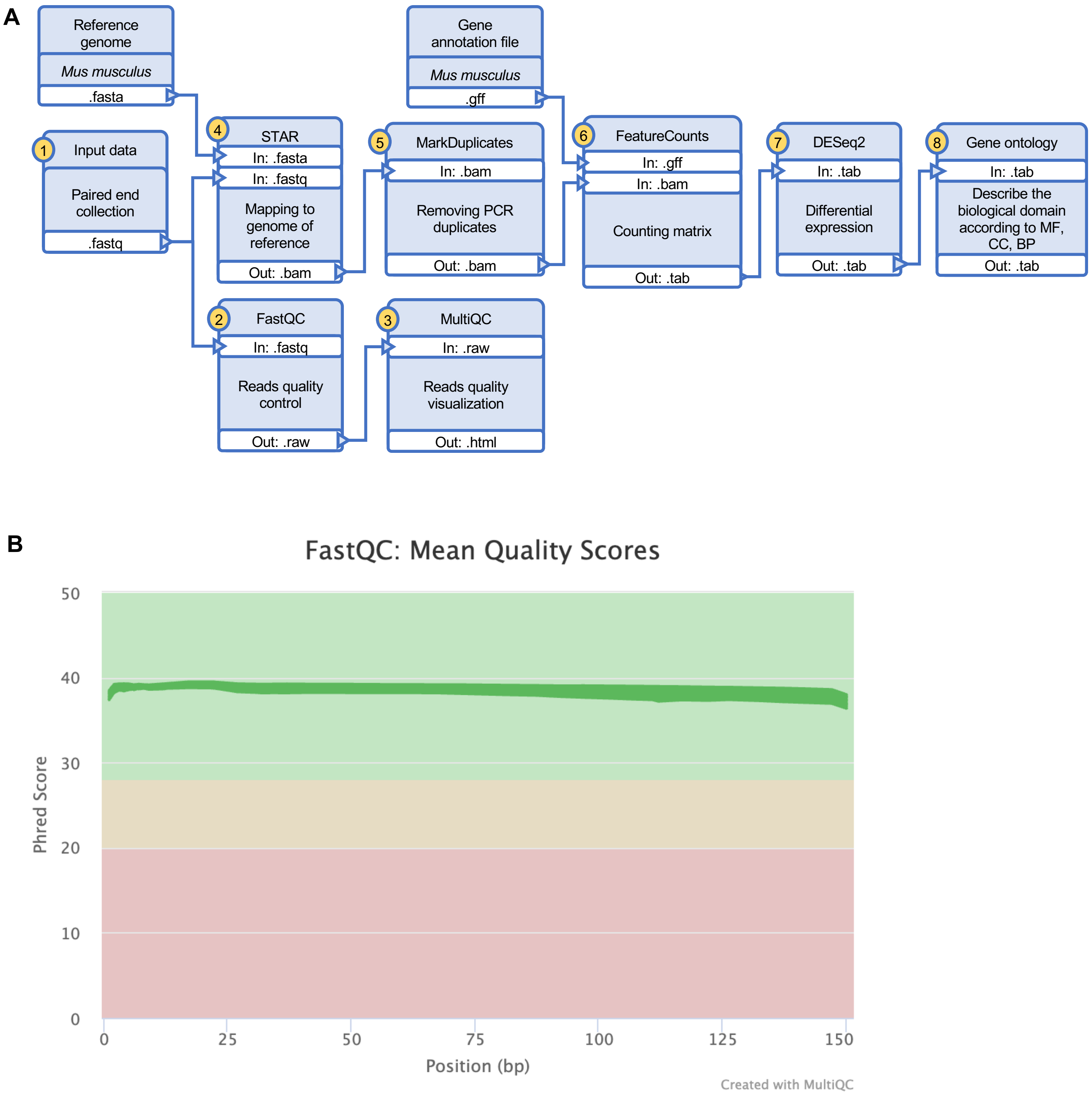

A Pituitary_ Mapping Summary

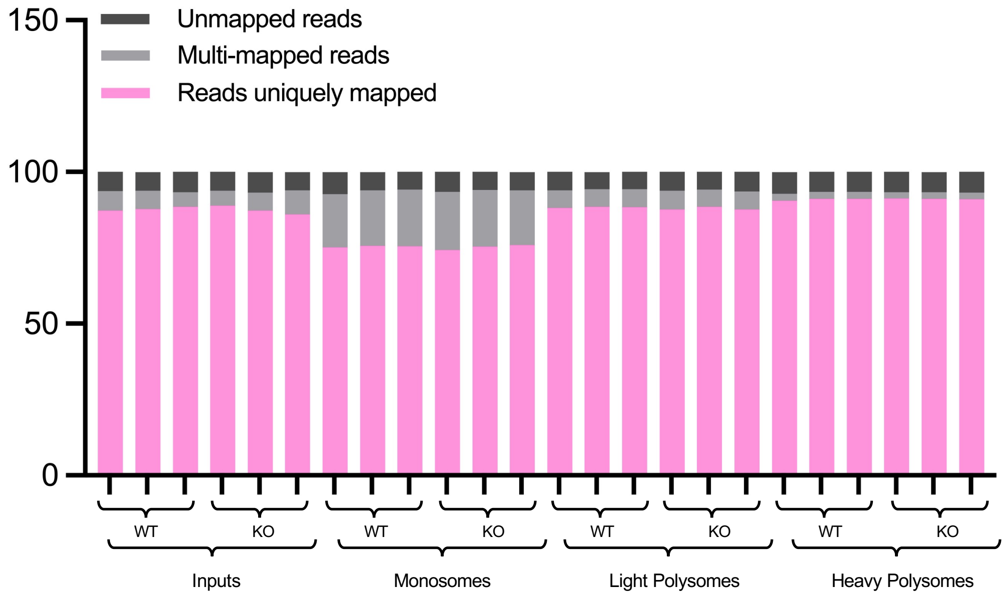


B Pituitary_ Counting Summary

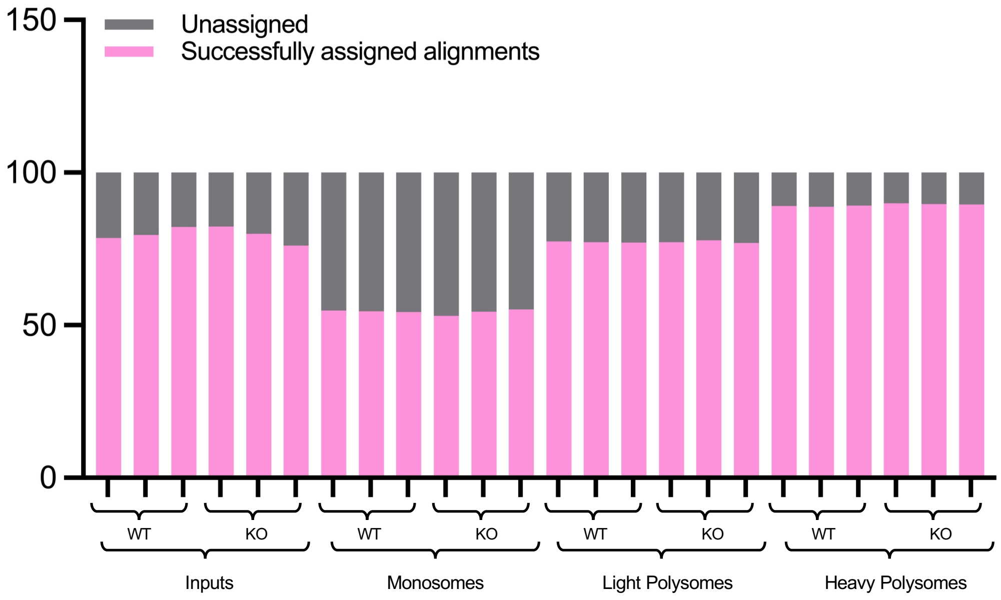


**Figure S3**

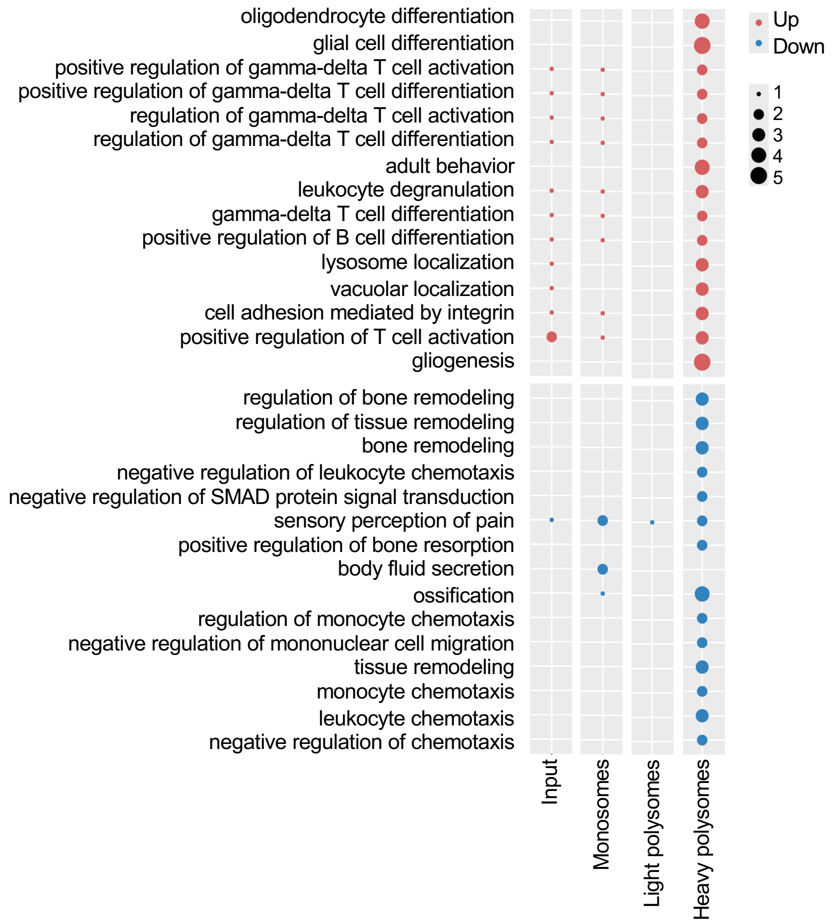

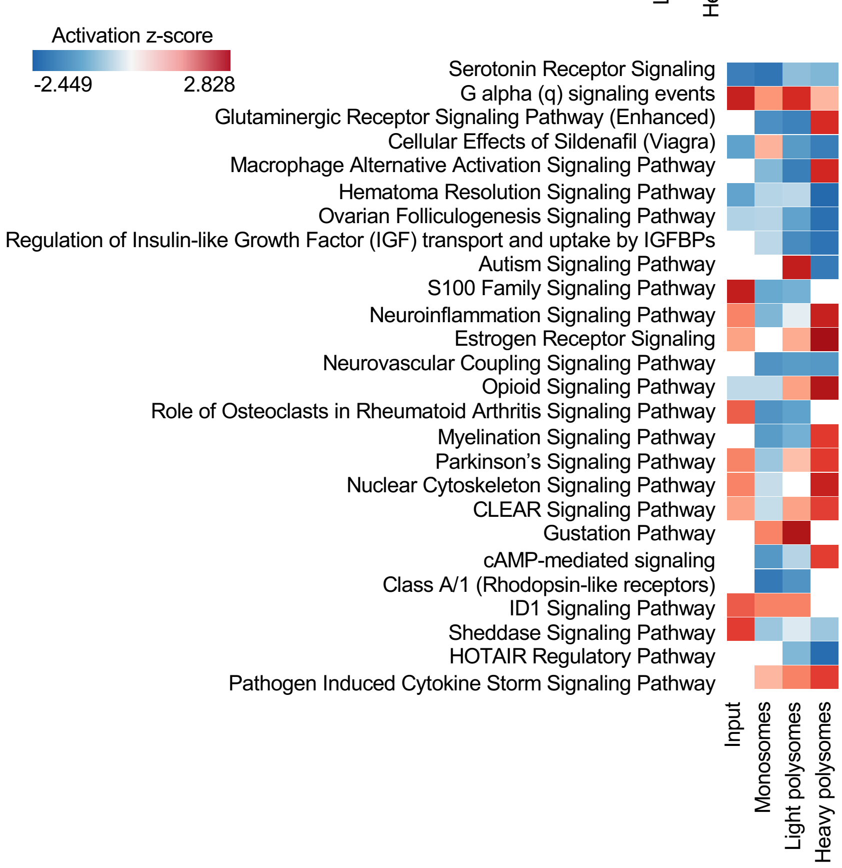

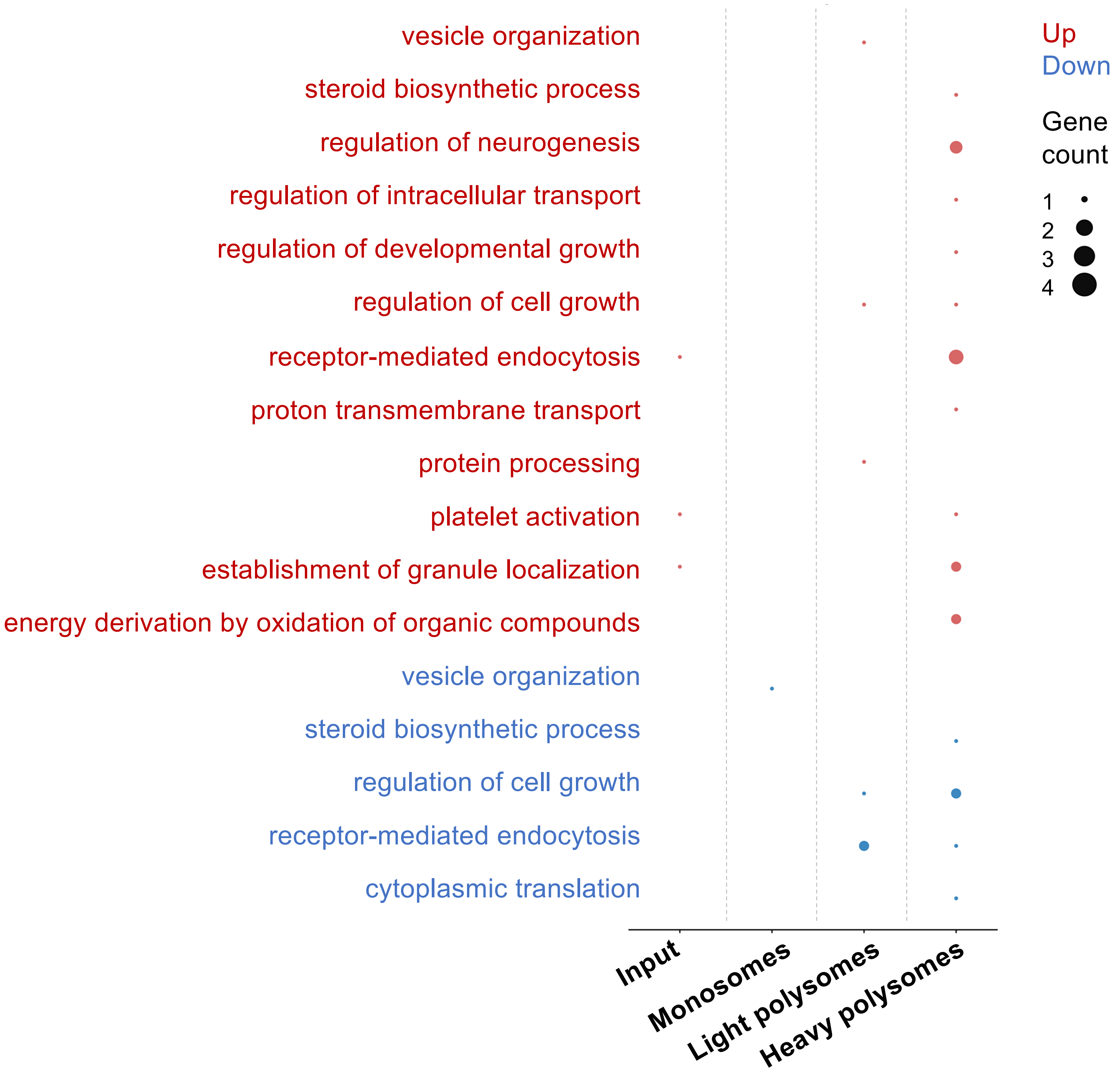
